# Modelling medulloblastoma pathogenesis and treatment in human cerebellar organoids

**DOI:** 10.1101/2024.08.14.607977

**Authors:** Thomas Willott, James G Nicholson, Yunchen Xiao, Olumide Ogunbiyi, Benjamin Draper, Talisa Mistry, Laura Donovan, Nicolae Radu Zabet, Ashirwad Merve, Sara Badodi, Silvia Marino

## Abstract

Faithful genetically engineered *in vivo* models of medulloblastoma (MB) are currently available only for some molecular subgroups, in keeping with recent studies showing the unique role of human-specific progenitors in the development of G3 and G4 MB subgroups. We generated human cerebellar organoids (CbO) from expanded potential stem cells (EPSC) and characterised their epigenetic and transcriptomic profile compared to the developing human cerebellum. We show the presence of sub compartment-specific cerebellar lineages linked to MB formation, including populations expressing signature genes of putative G3 and G4 MB cells-of-origin. We show that these lineages can be genetically engineered to model MB tumour onset. Moreover, we demonstrate that CbO sustain proliferation and invasion of G3/4 MB cells in a 3D co-culture model (CbO-MB) while preserving their molecular identity, and that treatment of CbO-MB with anti-tumour compounds recapitulate the efficacy of *in vivo* drug testing in xenograft models.

## Introduction

Medulloblastoma (MB) is a cerebellar tumour accounting for the largest proportion of brain cancer diagnoses in children. Molecular subgrouping based on transcriptomic, epigenomic and genomic features has collectively identified 4 MB subgroups: SHH (Sonic Hedgehog), WNT (Wingless-related Integration Site), Group 3 (G3) and Group 4 (G4), each further divided into subtypes with distinct prognosis and responses to therapy^1–4^. Despite these advances in classification, MB standard of care remains unchanged, with patients undergoing surgical resection followed by chemotherapy and/or radiotherapy^5^, which, albeit effective in a majority of patients, is associated with significant side effects.

The origins and molecular alterations for SHH and WNT MB are well characterised, leading to a surge in the development of molecularly targeted and/or immunotherapies in *in vitro* and *in vivo* models translating into early-phase clinical trials for patients in recent years^6^. The unclear origin and lack of universal pathway aberrations in G3/G4 MB subgroups along with the consequent lack of faithful pre-clinical models has hampered the identification of new treatment options for these tumours^7^. In particular, *in vitro* models for G4 MB, the most common MB subgroup, are highly underrepresented compared to other subgroups due to difficulties establishing 2D cell culture, whilst currently available G3 MB cell lines predominantly harbour *MYC* amplifications without encompassing the heterogeneity seen in these tumours^8,9^. The maintenance of primary MB cells as patient-derived orthotopic xenograft (PDX) models has overcome some of these issues^10–12^, however, the cost and resources required for these models limit their broader applications and scalability for drug testing.

MB formation is closely linked to deregulated cerebellar development^13^. Mapping lineage trajectories from MB single cell RNA sequencing (scRNA-Seq) data to human foetal cerebellar development has recently identified spatiotemporally restricted mitotic glutamatergic neuron progenitors in the rhombic lip subventricular zone (RL^SVZ^) harbouring photoreceptor (PRC) and unipolar brush cell (UBC) gene signatures as putative G3 and G4 MB cells of origin respectively^14,15^. These progenitors are halted in development prior to cell cycle exit leading to malignancy. Importantly, immunohistochemical and pseudo-time comparisons of relevant stages of human and murine cerebellar development showed significantly fewer of these proliferative progenitors in mice, raising the possibility that mouse models may not allow the study of key features of G3/G4 MB, including their onset^14,15^. Additionally, a human-specific transitional cell progenitor (TCP) population has been identified in the developing foetal cerebellum at 12-14 PCW, which is maintained or enriched in G3/4 MB tumours^16^. More recently, neural stem cells in the RL^VZ^ harbouring a *PRTG*^+^;*SOX2*^+^;*MYC*^high^;*NESTIN*^low^ signature have been linked to G3 MB onset^17^. These studies suggest that the failed maturation of early cerebellar cell lineages at specific sub-compartment location and developmental stage is linked to the ontogeny of G3/4 MB tumours.

Advances in organoid research have enabled the generation of new tools to study various aspects of cancer biology, including tumour initiation^18^, metastasis^19^ and as a co-culture model for drug testing^20^, with important implications for personalised medicine^21^. Methods to generate cerebellar organoids from induced pluripotent stem cells (iPSC) have been developed^22,23^ and characterised at single cell transcriptome level, achieving neuronal and glial differentiation with cerebellar sub-compartment identity^24–26^. In the context of MB modelling, current applications of cerebellar organoids are restricted to studies of neoplastic transformation in bulk genetically engineered organoids which lacked specificity for their respective cell-of-origin^27,28^, or in co-culture with established MB cell lines belonging to the SHH subgroup^29^.

Here, we have generated human cerebellar organoids (CbOs) from expanded potential stem cells (EPSC) and assessed their suitability to model G3/G4 MB. We have shown that they contain the putative MB cell-of-origin at stages of differentiation comparable to foetal cerebellar development and demonstrated that these cells can be genetically engineered to model early events in tumour initiation. In addition, we have analysed the suitability of CbOs as a co-culture model of G3/G4 MB providing insight into MB transcriptional state heterogeneity and tumour-microenvironment interactions. Finally, we show the value of the model as a pre-clinical drug testing tool.

## Results

### CbO display a cellular composition recapitulating human cerebellar development

We generated human CbO from two EPSC lines (CbO19 and CbO61) obtained from reprogramming of dura mater fibroblasts and extensively previously characterised^21,30^. Organoid differentiation was followed for 35 days, as outlined in previously published iPSC-derived protocols (Figure 1A and S1A)^22,23^. We observed a significant increase in organoid size throughout differentiation (Figure S1B) and the formation of characteristic flat-oval neural rosettes at Day 35 (D35) of culture (Figure 1B). Expression of the stemness marker *NANOG* showed a significant decrease by D7, confirming the commitment of the EPSC to differentiate (Figure S1C). Additionally, the neural stem cell (NSC) markers *NESTIN* and *SOX1* demonstrated early expression from D7-14, as well as the cerebellar homeobox and developmental genes *EN2* and *OTX2,* involved in early isthmic organiser signalling^62^ (Figure 1C and Figure S1D). Early progenitor markers *KIRREL2* and *PAX6* were expressed at D7-14 followed by markers of more mature precursors including *OLIG2/SKOR2* and *ATOH1/BARHL1* at D21-28, indicating the presence of GABAergic and glutamatergic lineages respectively (Figure 1C and Figure S1E, F). Interestingly, the transcription factor *EOMES,* expressed in glutamatergic progenitors and later in UBC, was expressed both in early (D7) and late (D28-35) stages of CbO maturation (Figure 1C and Figure S1F). The presence of GABAergic and glutamatergic granule neurons (GN) was confirmed by expression of CALBINDIN and Neu-N proteins respectively, proximal to SOX2-positive neural rosettes at D35 (Figure 1D and 1E). We observed similar patterns of expression throughout the differentiation of CbO19 and CbO61 (Figure 1C), in keeping with the conclusion that these CbOs reproducibly differentiate into cerebellar cell lineages.

**Figure 1.**
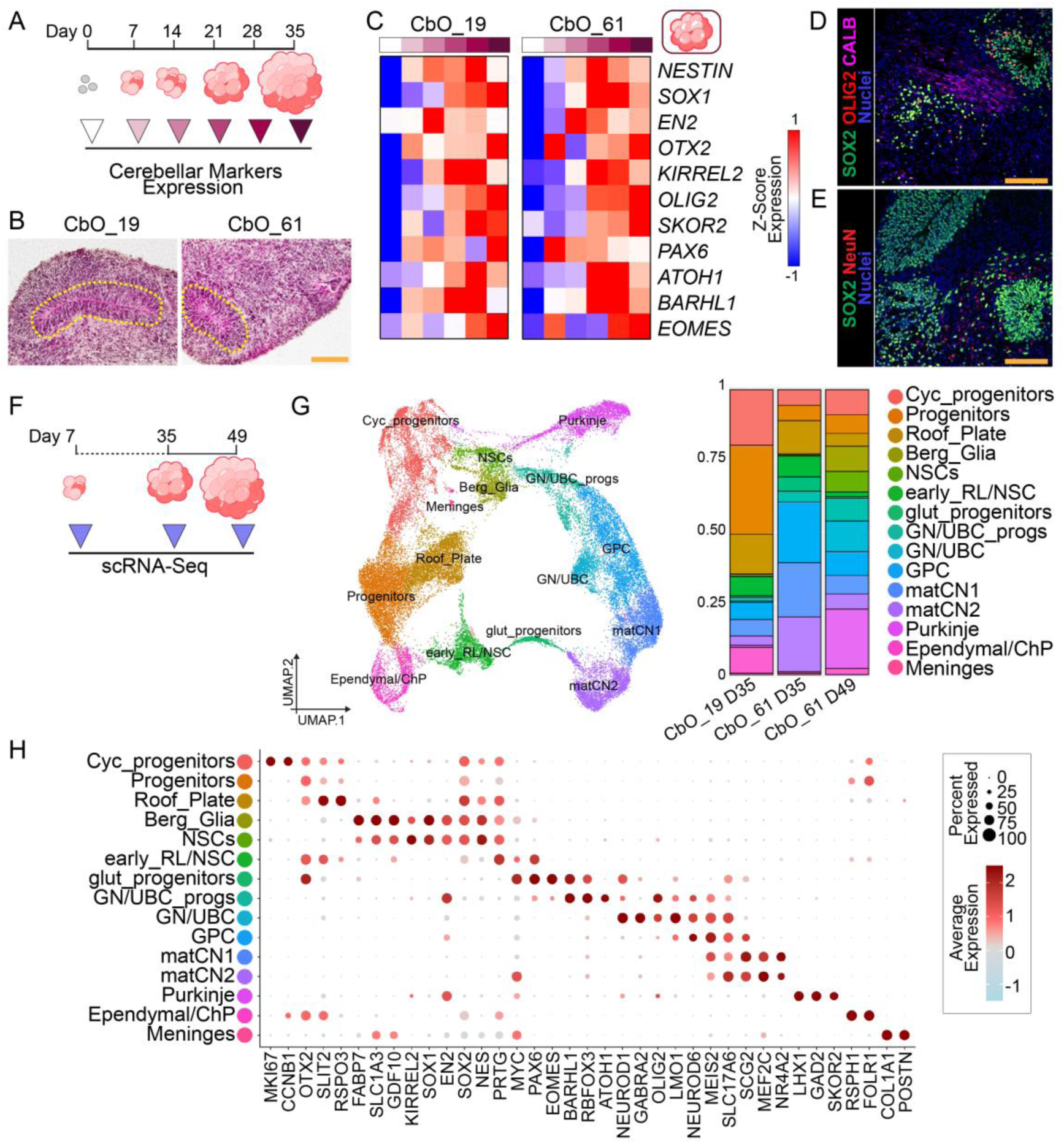
CbO is composed of diverse cerebellar lineages and displays characteristic structural organisation. A. Schematic of CbO samples used to assess expression of cerebellar markers. B. H&E staining of CbO_19 and CbO_61 showing formation of characteristic flat-oval neural rosettes within each structure. C. Heatmaps showing average expression of marker genes associated with cerebellar differentiation up to Day 35 (D35) of development for CbO_19 and CbO_61 derived from two independent EPSC lines (D0). Heatmaps show Z-scores for average dCT values from n = 3 independent batches of CbO. D. Representative immunofluorescence staining of OLIG2^+^ GABAergic progenitors (red), Calbindin^+^ (CALB^+^) mature neurons (pink) and SOX2^+^ neural stem and progenitor cells (green) in D35 CbO_61. E. Representative immunofluorescence staining of NeuN^+^ glutamatergic neuronal cells (red) and SOX2^+^ neural stem and progenitor cells (green) in D35 CbO_61. F. Schematic of CbO samples used in single cell RNA-Seq (scRNA-Seq) analysis. G. Uniform manifold approximation and projection (UMAP) plot of scRNA-Seq data from CbO_19 and CbO_61 at Day 35 and CbO_61 at Day 49 of differentiation (left) and bar plots showing percentages of annotated cell populations identified (right). H. Dot plot showing key marker genes used to identify major cell types in CbO (see also Figure S1M). Scale bars = 150 µm (B) and 100 µm (D and E).

To characterise in depth the cellular composition of CbO we performed scRNA-seq at D7, D35 and D49 timepoints to capture the earliest and more mature stages of cerebellar development and neurogenesis (Figure 1F). Initial data exploration revealed broad differences between D7 and the more heterogeneous D35 and D49 timepoints which were separated for downstream analysis (Fig S1G). Louvain clustering and Uniform Manifold Approximation and projections (UMAP) of D7 CbO cells identified three broad clusters corresponding to *PAX6*^+^ and *OTX2*^+^ glutamatergic progenitors, *PAX2*^+^, *PAX5*^+^ and *PKDCC*^+^ GABAergic progenitors and SOX9^+^ and S100B^+^ glial progenitors. We further identified a small population of nascent *NEUROD4*^+^ and *NEUROG1*^+^ glutamatergic neurons, suggesting an initial lineage commitment (Figure S1H and S1I). By D35 and D49 there was a greater degree of cellular diversity, and we identified 17 clusters (Figure S1J), with none specific to either timepoint (Figure S1K). Clusters were annotated based on their expression of marker genes (Figure 1G, H) and correlations to endogenous cell types from reference datasets of human developing cerebellum (Figure S1L)^16,42–44^. We identified non-committed *MKI67^+^* and *CCN1B^+^* cycling progenitors as well as *OTX2^+^* and *SOX2^+^* progenitor clusters. Two NSC clusters were identified, a broad cluster of *SOX1^+^, SOX2^+^, EN2^+^* and *NES^+^* NSCs and a more defined cluster expressing *SOX2^+^, PRTG^+^* and *PAX6^+^*. We found several clusters associated with glutamatergic lineages including two clusters of *BARHL1^+^* and *RBFOX3^+^* progenitors characterised by *PAX6* or *ATOH1* expression (glut_progenitors and GN/UBC_progs respectively). Additionally, we detected a *NEUROD1^+^* GN/UBC cluster also expressing mature glutamatergic neuronal markers (*LMO1* and *GABRA2*) and a *NEUROD6^+^* and *MEIS2^+^* glutamatergic progenitor cell (GPC) cluster. We further identified two mixed clusters of maturing cerebellar neurons (matCN1 and matCN2) positive for *MEIS2, SLC17A6* and *NR4A2* which are expressed by neurons located in the neural transitory zone (NTZ) during cerebellar development^44^. We further observed a neuronal cluster of GABAergic *SKOR2*^+^, *LHX1*^+^ and *GAD2*^+^ Purkinje cells as well as *FABP7^+^* and *SLC1A3^+^* Bergmann glial cells (Berg_Glia). In addition to cerebellar lineages, we also identified other hindbrain clusters containing cell types contributing to cerebellar development, including *FOLR1*^+^ and *RSPH1*^+^ ependymal/choroid plexus, *POST*^+^ and *COL1A1*^+^ meninges and *SLIT2*^+^ and *RSPO3*^+^ roof plate (Figure 1G, H).

Our data show that the cellular composition of CbO recapitulates the main cell types found during cerebellar development.

### Maturation of CbO follows conserved epigenetically-regulated central nervous system developmental pathways

Epigenetic regulation of gene expression, in particular DNA methylation, plays a fundamental role during central nervous system (CNS) development as it controls the spatio-temporal activity of cell lineage-specific genes^63^. Hence, we analysed the DNA methylation profile of CbOs during differentiation and maturation (Figure 2A). We identified genes showing the most variable time-dependent methylation status (VMGs) during CbO development and grouped them into time-dependent methylation programs by k-means clustering. Among genes exhibiting an increasing methylation status during CbO maturation (clusters 1 and 2) we found transcription factors linked to early cerebellum development including *PAX2*, *PAX5* and *PAX6* and to cell lineage specification such as *LMX1A* and *OTX2*. On the contrary, genes involved in neuronal differentiation and axon-guidance including *MAP2*, *MAP4*, *MAP7* and *GRIP1* were progressively hypomethylated during CbO differentiation (cluster 4 and 5) (Figure 2B). Gene Ontology (GO) pathway analysis of the identified VMGs revealed that early hypomethylated clusters were enriched for biological processes related to embryogenesis, fate specification, and organ development. Conversely, genes displaying later hypomethylation patterns were associated with neuronal differentiation, signal transduction, and structural functions such as synaptic junctions and neuron projections (Figure 2C). Analysis of human cerebellar samples (Figure S2A) revealed similar clustering of VMGs with distinct methylation patterns through human development (Figure S2B). GO analysis of human VMG clusters demonstrated shared earlier pathways with CbOs including those involved in proliferation and tissue development, followed by later pathways associated with signal transduction, neuron projection and catalytic activity (Figure S2C). Notably, we identified 1086 common VMGs demonstrating similar changes in methylation status between human and CbO samples (Figure 2D) suggesting that CbOs recapitulate epigenetic regulation of the human foetal cerebellum. Finally, we integrated DNA methylation and RNA expression analysis to identify genes with concordant methylation status and expression between D7 and D35 (i.e. hypomethylated and upregulated or hypermethylated and downregulated, Figure 2E). We found a significant enrichment for pathways involved in tissue morphogenesis, proliferation and receptor signalling for hypermethylated/downregulated genes expressed during early CbO development and enrichment of neuronal synapse pathways for hypomethylated/upregulated genes expressed later in CbO differentiation (Figure S2D).

**Figure 2.**
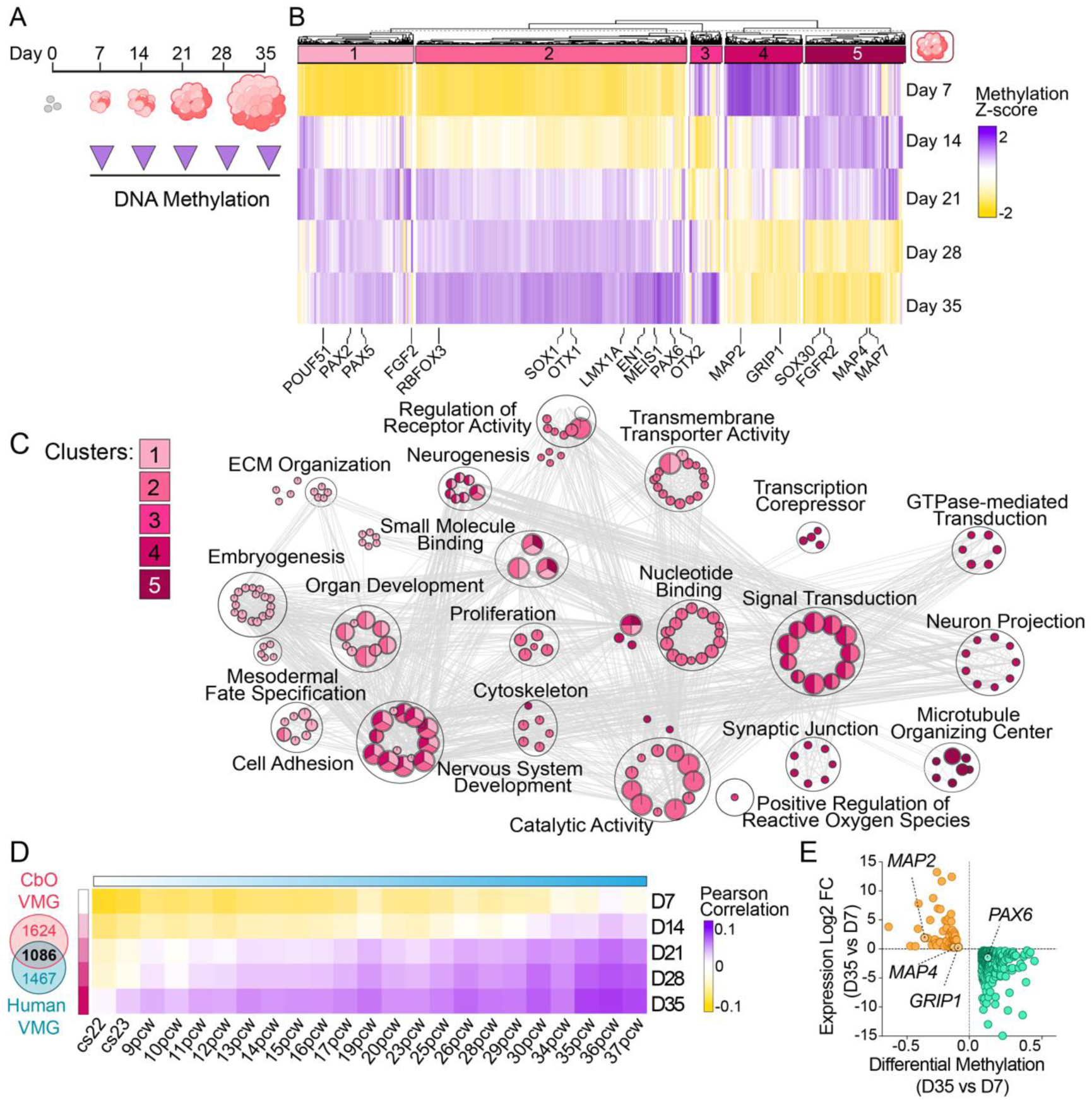
CbO development is epigenetically regulated similarly to human cerebellum. A. Schematic of CbO samples used to assess DNA methylation during development. B. Heatmap showing k-means clustering (n=5) of significant variable methylated genes across 35 days of CbO maturation. Mean beta values were taken for each timepoint from CbO_19 and CbO_61. Z-scores for average beta values of probes contained within each gene are plotted, with key cerebellar developmental and neuronal genes annotated. C. Bubble plot showing top 100 significant (adjusted p-value < 0.05) GO Biological Processes and Molecular Functions of genes from 5 clusters shown in **B**. Bubbles are coloured based on each cluster and size is proportional to number of genes in specific GO term. D. Venn diagram showing 1086 common variably methylated genes (VMGs) calculated through human (Carnegie stage 22 to 37 post conception weeks) and CbO (CbO_19 and CbO_61, Day 7 to 35) development (Upper). Pearson correlation of human and CbO samples on their beta values for 1086 common VMGs. E. Volcano plots of differentially expressed (DEG) and differentially methylated (DMG) genes between D35 and D7 of CbO maturation. Orange and green dots represent upregulated/hypomethylated and downregulated/hypermethylated genes respectively. Genes found in VMG clusters in B are highlighted.

Taken together, our data show that maturation of CbOs is governed by epigenetic regulation of key developmental pathways similar to the human cerebellum.

### CbOs reproduce the spatiotemporal cellular complexity required for medulloblastoma initiation

Next, we compared DNA methylation profiles of CbO and human foetal cerebellum samples to stage the development achieved. We observed the highest correlation of D35 CbO with 12-17 PCW cerebellum by comparing the methylation profiles of the top 1% of CpG probes with a linear correlation of methylation status across human foetal cerebellar development (Figure 3A and Figure S3A). Similarly, the application of an epigenetic clock algorithm trained on independent foetal brain datasets^57^ showed a positive correlation between differentiation of CbO and their predicted epigenetic age (R-squared = 0.8), confirming the staging of D35 CbO at 14 PCW and progressive maturation of the organoid (Figure S3B).

**Figure 3.**
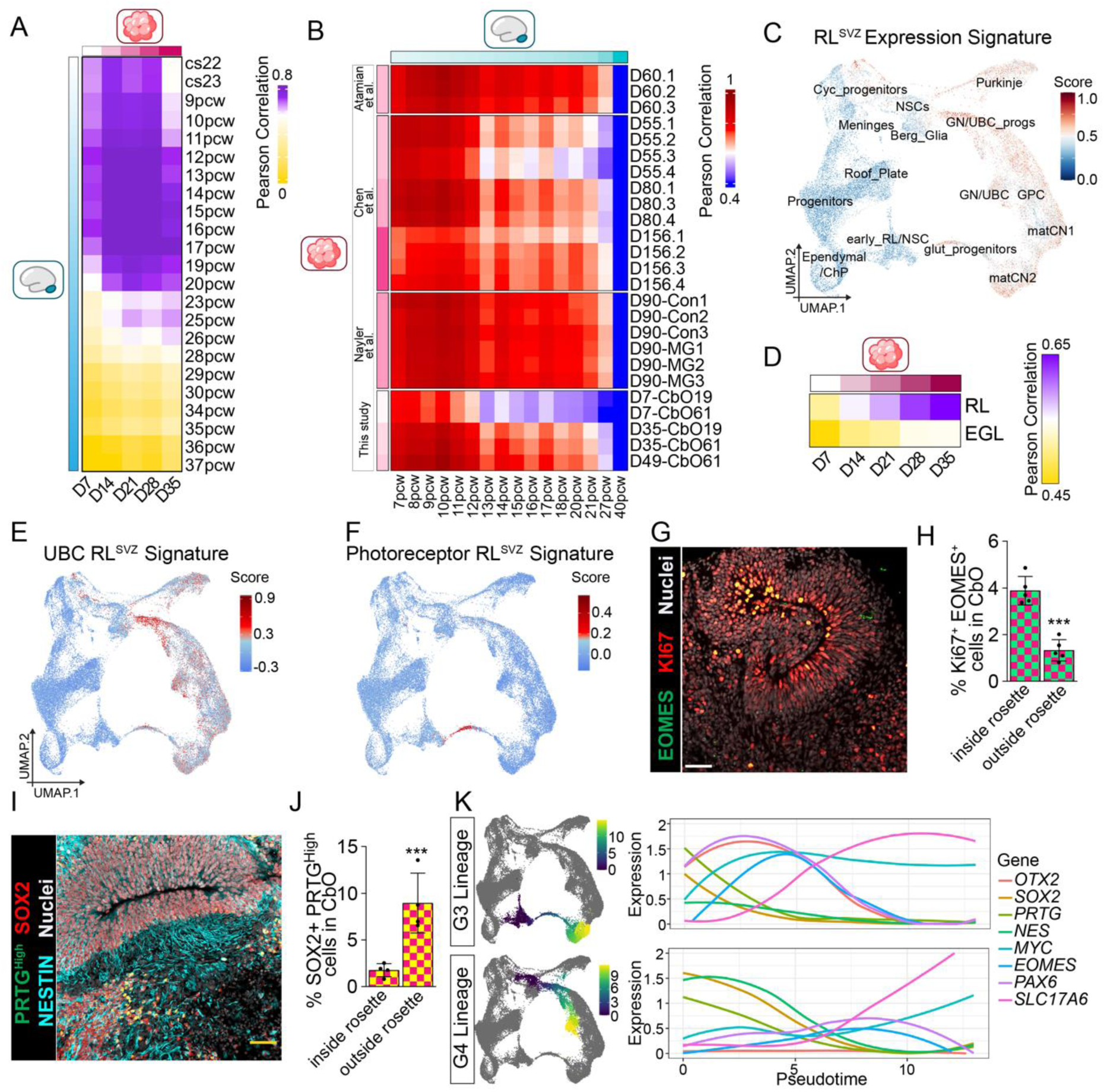
CbO displays spatiotemporal expression of G3/G4 MB cell-of-origin. A. Heatmap showing correlation of methylation status of top 1% (6886) age-dependent CpG probes between human developing cerebellum samples and CbO (see also Figure S3A). B. Heatmap showing correlation of top 1% (200) age-related genes between pseudo-bulk RNA-Seq of CbO and cerebellar organoids from previously published protocols^24–26^ and human foetal cerebellar samples^16,42–44^ calculated similarly to described above. C. UMAP plot of scRNA-Seq data from CbO at Day 35-49 of differentiation showing signature score for rhombic lip subventricular zone (RL^SVZ^) signature genes^15^. D. Heatmap showing correlation of average beta values of CbO to human microdissected rhombic lip (RL) and external granule layer (EGL) regions^15^ for differentially methylated probe set calculated between RL and EGL samples. E. UMAP plot of scRNA-Seq data from CbO at Day 35-49 of differentiation showing signature score for unipolar brush cell (UBC) RL^SVZ^ signature genes^15^. F. UMAP plot of scRNA-Seq data from CbO at Day 35-49 of differentiation showing signature score for photoreceptor RL^SVZ^ signature genes^15^. G. Representative immunohistochemistry pictures of D35 CbO_61 showing a neural rosette with cells positive for EOMES (green) and Ki67 (red). Nuclei are sown in grey. H. Quantification of staining shown in G of Ki67+EOMES+ nuclei with respect to CbO neural rosette structures. students t-test ^✱✱✱✱^p<0.0001. n = 5 regions of interest quantified. I. Representative immunohistochemistry pictures of D35 CbO_61 showing a neural rosette with cells positive for PRTG (green), SOX2 (red) and NESTIN (light blue). Nuclei are shown in grey. J. Quantification of staining shown in **I** of SOX2+PRTG^High^ nuclei with respect to CbO neural rosette structures. PRTG^High^ was defined as the top 50% of fluorescence signal for each image. students t-test ^✱✱✱✱^p<0.0001. n = 5 regions of interest quantified. K. Defined Group 3 (G3) and 4 (G4) medulloblastoma cell-of-origin trajectories selected for further analysis (G3 = early_RL/NSC, glut_progs, matCN2; G4 = NSCs, GN/UBC_progs, GN/UBC; see Figure 1H). Expression of genes associated with G3/4 medulloblastoma across each of the G3 and G4 trajectories.

Moreover, we analysed pseudo-bulk CbO RNA-seq data against the average expression of reference datasets^16,42–44^ from developing cerebellum and observed that D35 CbO show a high correlation in expression of age-related genes with 9-15 PCW (Figure 3B). Interestingly, our D35/49 CbOs acquire a similar expression profile to other cerebellar organoids cultured for longer time periods^24,25,43^ (Figure 3B).

Putative G3/4 MB cells-of-origin were first described in the RL^SVZ^, and we indeed saw an enrichment of an RL^SVZ^ gene signature score in the glutamatergic cerebellar neuronal and progenitor clusters, which was depleted in less differentiated cycling progenitors and NSC (Figure 3C). Correlation of CbO methylation levels to microdissected RL and EGL samples^14^ revealed an increased correlation through CbO development with both regions, reflecting the expanding populations of predominantly glutamatergic neuronal progenitors in the CbO, and showing high correlation of D35 CbO specifically with RL (Figure 3D).

Previous studies showed enrichment of photoreceptor and UBC expression signatures in RL^SVZ^ (PRC.RL^SVZ^ or UBC.RL^SVZ^), with a high expression in G3 and G4 MB putative cells-of-origin respectively. Quantification of these signatures within CbOs confirmed their enrichment in discrete clusters of glut_progenitors or GN/UBC_progs respectively, with the UBC.RL^SVZ^-enriched GN/UBC_prog cluster also highly correlated to TCP cells (Figure S1L), a progenitor population previously associated with G3/4 MB^16^. Importantly, both of these progenitor clusters were characterised by *EOMES* expression (Figure 3E, F and Figure S3C) and we observed EOMES^+^Ki67^+^ cells located predominantly within the neural rosettes in CbO, reminiscent of the human-specific G3 and G4 MB-susceptible progenitors (Figure 3G, H).

Notably, the early_RL/NSC cluster, which RNA velocity analysis predicts to give rise to PRC.RL^SVZ^-enriched glut_progenitors, presents high expression of *PRTG, SOX2* and *MYC* and low expression of *NESTIN* (Figure 1H, S3D), mirroring the gene signature found in a *PRTG*^+^;*SOX2*^+^;*MYC*^high^;*NESTIN*^low^ stem cell population in the RL^VZ^ recently proposed as a putative G3 cells-of-origin^17^. Immunohistochemical staining revealed PRTG^High^;SOX2^+^ cells outside of CbO neural rosettes in areas presenting concomitant low expression of NESTIN (Figure 3I, J), suggesting a spatial complexity to glutamatergic NSC/progenitor cells in CbOs with differing MB cell-of-origin signatures.

Given that pre-malignant G3/4 MB cells-of-origin are thought to persist in the RL, likely due to their stalled differentiation^15^, we performed pseudotime and RNA velocity analysis through the D35/49 UMAP (Figure S3D,E). Plotting G3 and G4 trajectories through each UMAP demonstrated sequential expression of glutamatergic neuronal genes associated with early lineage commitment, progenitor expansion and more mature neurons (Figure S3F). We observed an earlier high expression of the G3-associated genes *PRTG, OTX2* and *MYC* during sustained *SOX2* expression in the G3 trajectory in line with G3 cells-of-origin harbouring stem and early progenitor phenotypes (Figure 3K). Whereas, in the G4 trajectory, MB-associated genes *EOMES* and *PAX6* exhibited high expression relatively later in pseudotime, after *SOX2* loss and before upregulation of the mature glutamatergic marker *SLC17A6*, highlighting a more committed progenitor cell population (Figure 3K).

Taken together, our data show that D35 CbOs display an epigenetic and transcriptional landscape mirroring the cerebellar developmental stage and cellular profiles of G3/G4 MB initiation.

### Genetic editing of MB cell-of-origin populations in CbO models tumour onset

Leveraging the observation that CbO mirrors the cellular complexity of the human cerebellum at the time of G3/G4 MB formation, we sought to model tumour initiation in CbOs by specifically targeting their relevant cell-of-origin.

We set out to overexpress the GFP-tagged *c-MYC* oncogene in G3/G4 progenitors and to achieve selective spatio-temporal targeting of the cells we used a Tet-inducible system driven by the promoter of *PAX6* or *EOMES*, transcription factors known to have a role in MB^64,65^ and expressed in MB cell-of-origin clusters at D35 of CbO differentiation (Figure 1H, S3C, 3K). EPSCs transduced with either *pEOMES;MYC-GFP* or *pPAX6;MYC-GFP* were used to generate CbOs and c-*MYC* overexpression was induced at D35 of maturation (Figure 4A). A *BFP* tag was added to p*EOMES* and p*PAX6* constructs to allow purification of the engineered cells from non-induced CbOs and we confirmed retention of early stem cell markers expression in transduced EPSC (Figure S4A). Foci of c-MYC-GFP^+^ cells were observed as early as 4 days after doxycycline treatment in both editing systems, with their expansion observed upon maturation (Figure S4B). D63 CbO showed clusters of highly proliferative GFP^+^Ki67^+^ cells, which corresponded to areas of densely packed hyperchromatic and mitotically active cells (Figure 4B).

**Figure 4.**
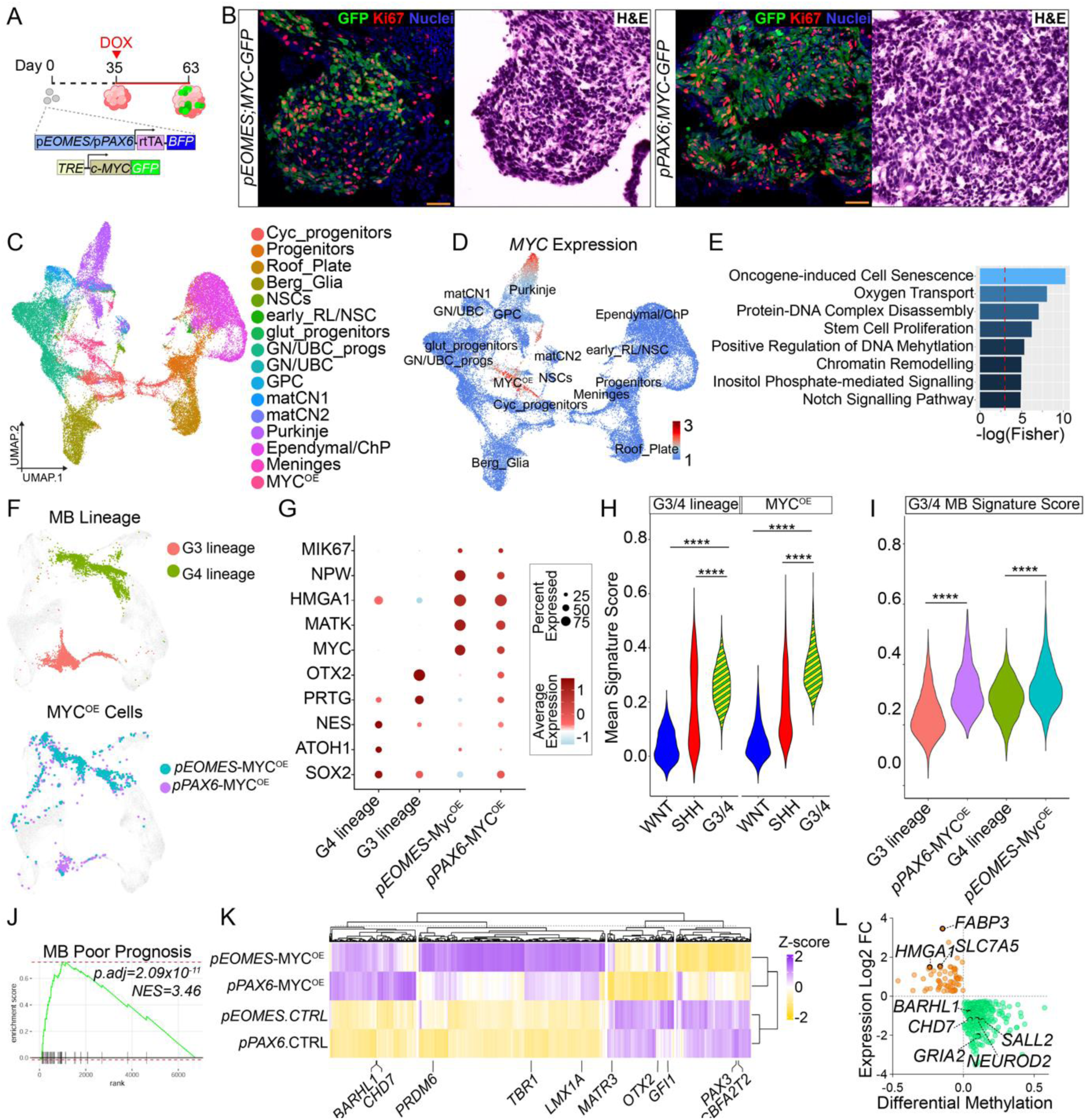
Edited medulloblastoma lineages-of-origin display features of MB dysregulation upon MYC overexpression. A. Schematic of cerebellar organoid (CbO) editing strategy (see Methods). B. Representative immunohistochemistry staining for GFP (green) and Ki67 (red) (left) and haematoxylin and eosin staining (right) for CbO_19 collected at Day 63 of differentiation for dox-induced *pEOMES;MYC-GFP* and *pPAX6;MYC-GFP* organoids. C. UMAP plot of scRNA-Seq data from *pEOMES;MYC-GFP, pPAX6;MYC-GFP* and control CbOs at Day 63 of differentiation and colour by cell type. D. UMAP shown in D with *MYC* expression plotted. E. Gene ontology enrichment bar plot of MYC^OE^ cluster marker genes. F. UMAP plot of control D35/40 CbO scRNA-Seq data highlighting Group3 (red) and Group4 (green) MB lineages of origins (Top). Projection of MYC^OE^ cells from D63 *pEOMES;MYC-*GFP (blue) *and pPAX6;MYC-GFP* (purple) CbOs onto control UMAP embeddings. G. Dot plot showing G3 and G4 lineage-of-origin marker and MB-associated genes in G3/4 lineage genes compared to MYC^OE^ cells from D63 *pEOMES;MYC-GFP and pPAX6;MYC-GFP* CbOs. H-I. Violin plot showing mean signature scores for WNT, SHH or G3/G4 MB subgroups^45^ in G3/4 lineages-of-origin and MYC^OE^ cells from D63 *pEOMES;MYC-GFP and pPAX6;MYC-GFP* cells CbOs. J. Gene set enrichment plot of the POMEROY_MEDULLOBLASTOMA_PROGNOSIS_DN gene set in the MYC^OE^ cell compared to G3/4 lineages-of-origin, NES=3.46, padj=2.09E-11 K. Heatmap showing z-scored average beta values of probes contained within differentially methylated genes between MYC^OE^ (*pEOMES;MYC-GFP, pPAX6;MYC-GFP)* and control medulloblastoma lineage-of-origin samples (*pEOMES.*ctrl, *pPAX6.*ctrl). Key cerebellar and medulloblastoma associated genes are annotated. L. Scatter plot of differentially expressed (DEG) and differentially methylated (DMG) genes between MYC^OE^ cells and G3/4 cell-of-origin. Orange and green dots represent upregulated/hypomethylated and downregulated/hypermethylated genes respectively.

To investigate the transcriptional consequences of *c-MYC* activation in the putative G3/4 cells-of-origin, we next performed scRNAseq on edited CbOs, transferring cell type labels from our previously annotated D35/49 dataset. c-*MYC* expression was particularly high in a cluster made up almost exclusively of cells from *pEOMES;MYC-GFP* and *pPAX6;MYC-GFP* CbO samples (cluster 12) that we therefore labelled as *MYC* overexpressing (MYC^OE^) cluster (Figure 4C, D and Figure S4C, D). Gene signature scoring confirmed the proliferative status of the MYC^OE^ cluster as well as its similarity to G3 MB^47^ (Figure S4E, F). Gene set enrichment analysis of the MYC^OE^ cluster confirmed enrichment of downstream MYC targets (Figure S4G) and GO enrichment of upregulated cluster markers highlighted terms consistent with epigenetic dysregulation (protein-DNA complex disassembly, chromatin remodelling, regulation of DNA methylation) and signalling pathways linked to MB (Notch signalling pathway, inositol phosphate-mediated signalling) (Figure 4E)^35,66^. Interestingly, we found enrichment of oncogene-induced senescence, a result in apparent contrast with the relatively proliferative signature scoring for this cluster but likely reflective of the countervailing forces of tumour suppression and oncogenic signalling as these cells undergo transformation.

Transformation of MB cells-of-origin has been linked to their stalled differentiation^15^, we therefore projected MYC^OE^ cells onto the D39/45 CbO UMAP (Fig 4F). As expected, the MYC^OE^ cluster from *pEOMES;MYC-GFP* CbO samples (p*EOMES-* MYC^OE^) overlaid the G4 lineage whereas those from the *pPAX6;MYC-GFP* CbOs (p*PAX6*-MYC^OE^) were more equally distributed between the G4 and G3 lineages. Notably, G4 lineage and p*EOMES-* MYC^OE^ cells express *ATOH1*, while the G3 lineage and p*PAX6*-MYC^OE^ express higher levels of *PRTG* and *OTX2* (Fig 4G), supporting our choice of *PAX6*/*EOMES* promoters to target specific progenitor clusters.

We next scored cells for patient-derived MB gene signatures, confirming that both our G3/4 lineage and MYC^OE^ clusters were more similar to G3/4 subgroup (Figure 4H). Importantly, we observed a significantly higher G3/4 MB signature score (Figure 4I) in engineered MYC^OE^ cells compared to their G3/4 lineage cells-of-origin alongside with significant depletion of gene sets associated with neuronal differentiation and enrichment of a gene set corresponding to poor MB prognosis (Figure 4J and S4H).

We further investigated the methylation profiles of sorted GFP^+^-MYC^OE^ cells as compared to BFP^+^ lineage-of-origin controls and found that they clustered with G3 MB patient tumours, as assessed by a MB subgroup-specific CpG probe set^1,59^ (Figure S4I). Furthermore, when focusing the analysis on DMPs between MYC^OE^ and lineage-of-origin we observed a stronger correlation to G3 MBs for the *pPAX6-*MYC^OE^ compared to *pEOMES-*MYC^OE^ model (Figure S4J). Interestingly, analysis of DMGs revealed model-specific deregulation of genes with *pPAX6-*MYC^OE^ demonstrating specific hypomethylation of *OTX2* and *GFI1*, known drivers of G3 MB onset^27^, whilst *pEOMES-*MYC^OE^ showed hypomethylation of genes with mutually exclusive roles to *OTX2* in G3/4 MB pathogenesis and cell fate decisions including *PAX3* and *CBFA2T2*^15,64^ (Figure 4K). Integration of DNA methylation and gene expression data further showed a concordant positive regulation of several key genes (*HMGA1*, *SLC7A5* and *FABP3*) and negative regulation of genes associated with neuronal differentiation (*SALL2*, *NEUROD2* and *GRIA2*) (Figure 4L).

Taken together, we show that using CbO we can model features of MB onset in G3/4 lineages-of-origin through *c-MYC* overexpression including increased proliferation, stalled development and transcriptional changes consistent with epigenetic dysregulation and poor MB outcome.

### CbOs sustain co-cultured medulloblastoma cells proliferation and invasion while preserving their molecular identity

As well as utilising CbO to model early events in MB cell-of-origin transformation, we also set out to assess whether CbO could provide a physiologically relevant 3D environment for sustaining growth of human MB cells. We co-cultured GFP-tagged G3/4 MB cell lines (CHLA-01-Med, CHLA-01R-Med and ICb1299) as well as G3 MB primary cells (Med211-FH) maintained as a PDX with D35 CbO when cerebellar lineages are clearly defined (Figure 5A). All MB lines successfully attached and infiltrated the CbO with considerable cell growth after 14 days of co-culture (Figure 5A and S5A-C). Morphological analysis of CbO bearing MB cells (CbO-MB) showed a highly cellular and proliferative pleomorphic embryonic tumour, in keeping with MB, diffusely infiltrating the adjacent CbO tissue (Figure S5D-F). We observed positivity for the proliferation marker Ki67 in around 30% of GFP^+^ MB cells (Figure 5B), thus confirming that CbOs can sustain MB cells proliferation and infiltration.

**Figure 5.**
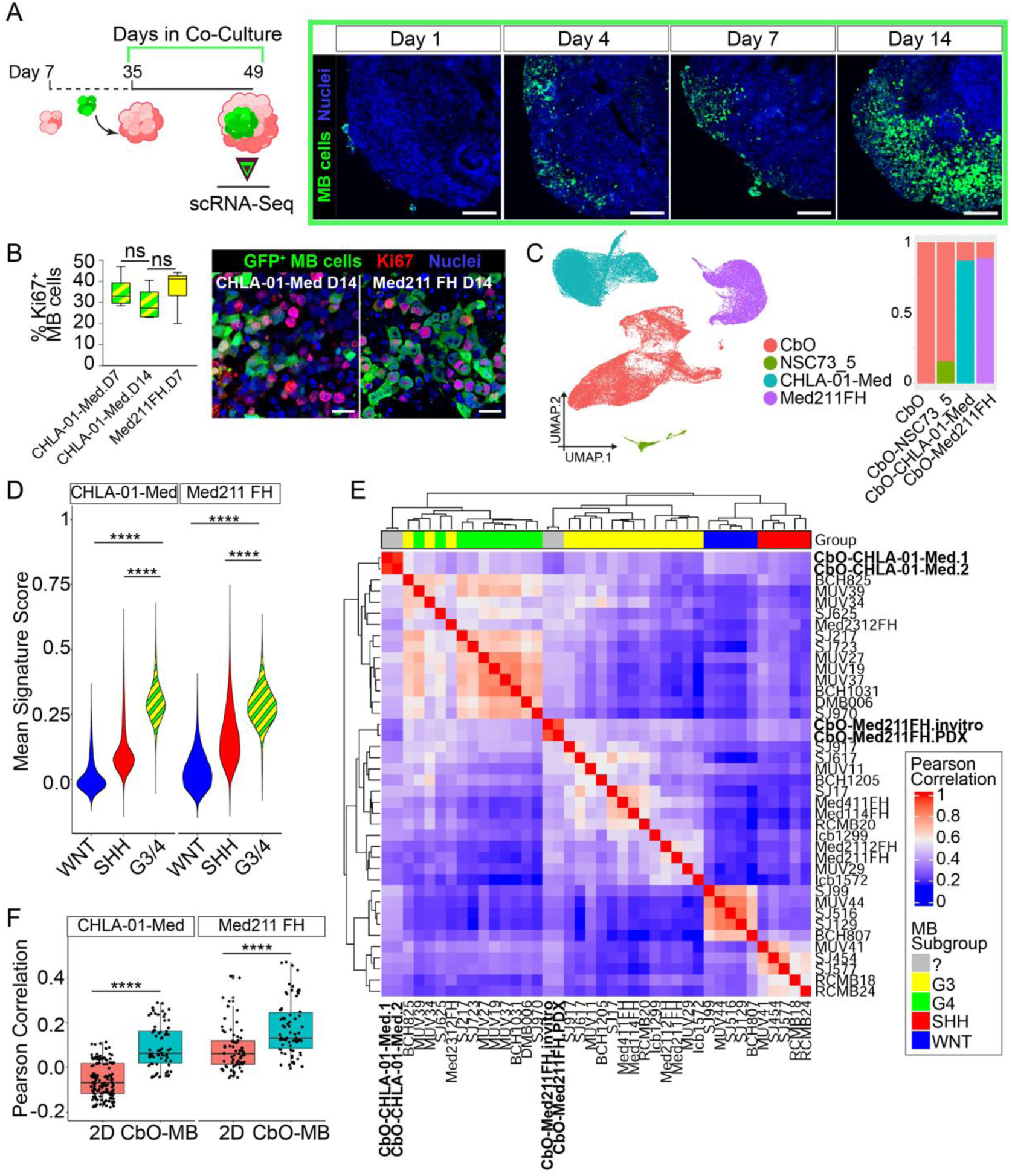
CbO enables MB growth whilst also retaining MB transcriptional profiles closer to those of patient tumour samples. A. Schematic of protocol used to co-culture cerebellar organoids (CbO) with medulloblastoma (MB) showing CbO-MB sample used for scRNA-Seq (left panel). Representative immunofluorescence staining of GFP^+^ CHLA-01-Med MB cells upon one, four, seven and fourteen days of co-culture with CbO_61, scale bars = 100 µm (right panels). B. Box and whisker plot showing percentage of MB cells (GFP^+^) expressing the proliferative markers Ki67 upon 7 and 14 days of co-culture in CHLA-01-Med and Med211FH cells. Graphs report mean ± SEM, unpaired t-test, n = 6 independent CbO-MB analysed, ns = not significant (left). Representative immunofluorescence staining of GFP^+^ and KI67^+^ MB cells upon 14 days of co-culture with CbO, scale bars = 100 µm (right). C. UMAP plot of scRNA-Seq data from D49 control CbO_61, CbO-Med211-FH, CbO-CHLA-01-Med and control CbO-NSC73_5 co-cultures, coloured by genotype (left). Barplot showing proportion of cells from each genotype in the different experimental models (right). D. Violin plot showing signature scores for WNT, SHH or G3/G4 MB subgroups^45^ in CbO-MB. One-way ANOVA ^✱✱✱✱^p<0.0001. E. Heatmap showing correlation scores between expression of pseudo-bulk RNA-Seq of MB fractions only from CbO-Med211-FH and CbO-CHLA-01-Med and reference MB samples. Clustering based on expression profile has been used to assess molecular subgroup of CbO-MB models. F. Boxplots plot showing correlations between same MB cell line grown as 2D (red) or 3D (blue) model and MB samples shown in D. One-way ANOVA ^✱✱✱✱^p<0.0001.

To explore the effect of MB cells on CbO, we performed scRNA-Seq on pooled CbO-MB samples (CHLA-01-Med or Med211-FH) after 14 days of co-culture (Figure 5A), as well as control CbOs co-cultured with cerebellar NSCs (NSC73.5, CbO-NSC). Louvain clustering and inferred copy number analysis identified CHLA-01-Med and Med211-FH MB cells (Fig 5C, S5G-H) expressing tumour-specific markers (*SIX3*/*APOE* and *NEUROD1*/*BCAT1* respectively) as well as shared MB-specific markers we previously observed in our modelled MYC^OE^ cells (*HMGA1*, *MYC*, *YBX3* and *TRAP1*). NSC73.5 cells could be distinguished by their separation in the UMAP, and expression of stemness markers (*SOX2/9*), doublecortin (*DCX*) and transcription factors governing interneuron development (*DLX1/2*) (Figure S5I). In keeping with the highly proliferative properties of tumour cells, we identified MB cells representing more than 80% of the sequenced CbO-MB cells as compared to NSC73.5 forming around 20% of CbO-NSC (Figure 5C).

To investigate the degree to which co-cultured CbO-MB cells retain their subtype specificity, we computationally isolated co-cultured MB cells and scored them for patient-derived MB gene signatures^45^. We observed a significant increase in average G3/G4 metaprogram scores when compared to WNT or SHH scores (Figure 5D), indicating that both MB cell lines retained their molecular subtype in the CbO-MB model. We next correlated pseudobulk transcriptional profiles with the same reference cohort of patient MB tumours^45^. As expected, CHLA-01-Med fell within an intermediate subcluster containing both G3 and G4 primary tumours, whilst Med211-FH clustered with G3 tumours only (Figure 5E). Label transfer of G3 and G4 tumour scRNA-seq profiles onto co-cultured MB clusters further confirmed these subgroup identities (Figure S5J). Studies of glioblastoma have shown that tumour cells more closely recapitulate their parental tumours when co-cultured in brain organoids as compared to 2D models^21^. We therefore correlated expression profiles of each MB line co-cultured with CbO or in 2D with patient tumour samples^35,45^. Importantly, we observed a significantly higher correlation in all co-cultured MB cells compared to their 2D cultured counterparts (Figure 5F), suggesting that CbO-MB better phenocopy *in vivo* primary tumours.

These data show that CbO represents a novel 3D model for G3 and G4 MB cell growth and expansion which is superior at preserving features of patients’ tumour samples as compared to 2D cultures.

### Group 3 MB cells harbour a subset of cells with myogenic differentiation

We next analysed the MB fraction of CbO-MB co-cultures, performing Louvain clustering and correlating cluster marker signature to each other and to references of the developing cerebellum (Figure 6A-C and Figure S5K). In line with previous studies, both cell lines were predominantly composed of clusters corresponding to cycling and non-cycling GCP-like cells (clusters 1,7, and 2 for Med211-FH; clusters 3 and 0 for CHLA-01-Med). Overall, Med211-FH clusters correlated more closely with developing cerebellar cell types than those from the CHLA-01-Med cell line and expressed genes consistent with a more neural phenotype (*SYT1, NEUROD1*). We also identified two pairs of smaller cell states that were common to both MB lines, clusters 8 and 5 and clusters 4 and 6 from Med211-FH and CHLA-01-Med, respectively (Figure 6D). Clusters 8 and 5 correlated to glial progenitor/NSC developing cerebellar cell types (Figure S5K) and specifically expressed stemness markers (*SOX2*, *NTRK2*) (Figure 6C). GO enrichment of the 44 shared marker genes highlighted terms associated with neuronal development and radial glia differentiation (Figure 6E). Clusters 6 and 4 showed a phenotype more consistent with immune reference cells and expressing immune-associated markers such as *IL17B* (Figure 6C and Figure S5K). Further analysis of the 127 shared marker genes between clusters 6 and 4 revealed a strong enrichment of GO terms associated with myogenic development and differentiation (Figure 6F). This included expression of key transcription factors governing myogenic differentiation (*MYOG, MYOD1* and *HLHL41*) as well as components of the myosin light chain (*MYL4*) and troponins (*TNNT1*) (Figure 6C) with MYOGENIN^+^ MB cells confirmed by IHC in CbO-MB co-cultures (Figure 6G). Previous histological studies have reported aberrant myogenic differentiation of MB cells^67,68^, but the transcriptional state associated with such phenotypes has not previously been described. By scoring reference MB samples^45^ for our CbO-MB derived myogenic state signature, we observed the highest score in the WNT MB subtype (Figure 6H) while analysis of Group 3 MB revealed a subset of cells scored highly for our signature suggesting heterogenous expression of this metaprogramme (Figure 6H). Indeed, upon sub-setting, and re-clustering Group 3 MB samples we observed two clusters, 4 and 6, which expressed this myogenic signature strongly (Figure 6I). These myogenic cells were found in 11/16 samples ranging from 0.3-70% of total cells (Figure 6J). Scoring of the myogenic signature in a larger cohort of 763 MB samples^1^ revealed a similar distribution across MB groups (Figure 6K), with an enrichment in the poor prognosis MYC-amplified Group3_gamma subtype (Figure 6L) and a significant reduction of survival in samples scoring highly (> median) for the myogenic signature (Figure 6M).

**Figure 6.**
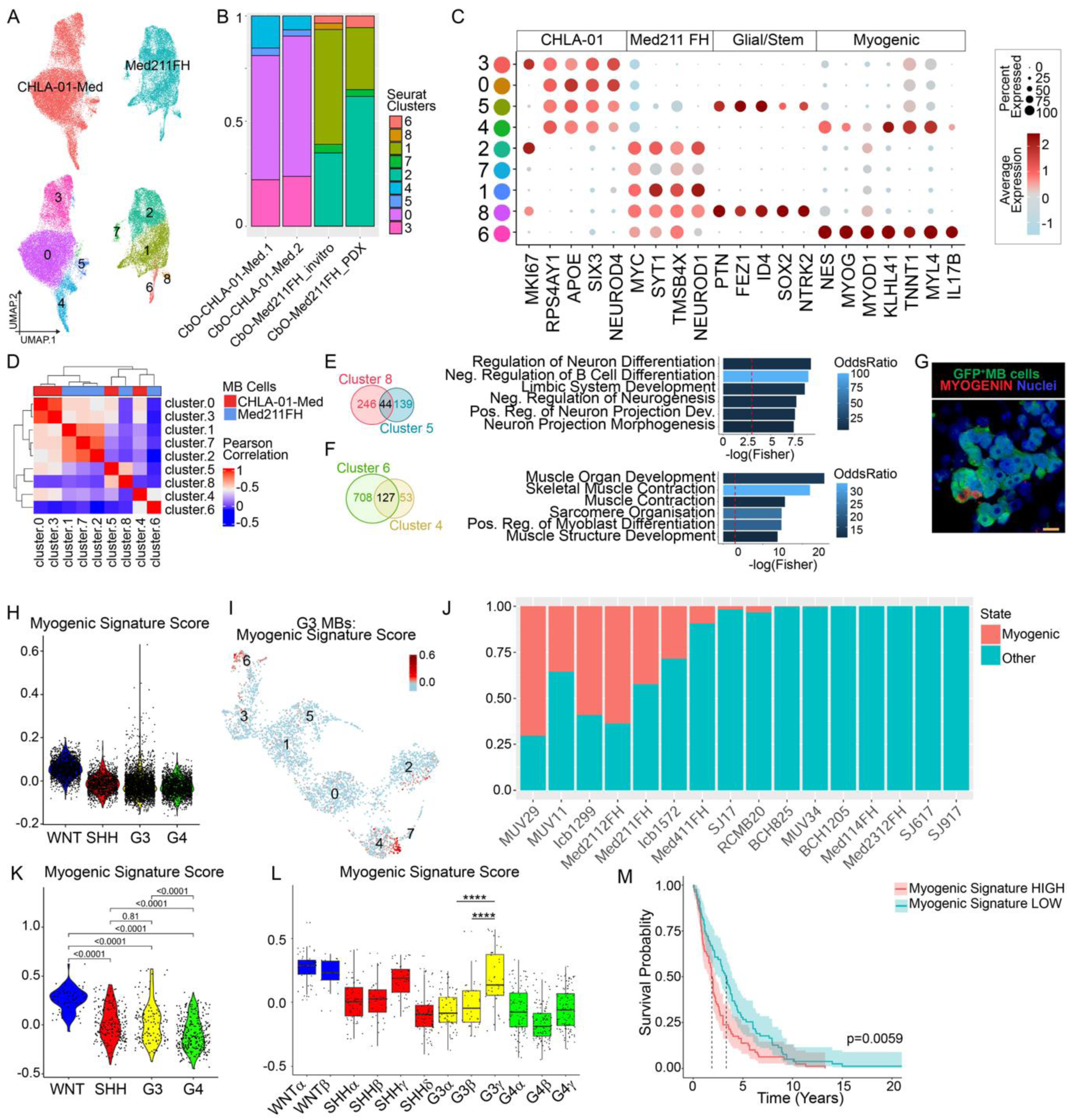
CbO-MB uncovers a subset of Group3 MB cells with a myogenic transcriptional state. A. UMAP plot of scRNA-Seq data from MB cells only from D49 CbO-Med211-FH and CbO-CHLA-01-Med CbO-MB co-cultures, coloured by genotype (top), and Louvain clusters (bottom). B. Barplot showing proportion of MB cells from each CbO-MB co-culture in each cluster. C. Dot plot showing key marker genes used to identify MB cell types in MB cells only from D49 CbO-Med211-FH and CbO-CHLA-01-Med CbO-MB co-cultures. D. Heatmap showing Pearson correlation of MB cell clusters across the top 250 specific markers for each cluster. E. Venn diagram showing the intersect of CbO-Med211-FH cluster 8 and CbO-CHLA-01-Med cluster 5 marker genes (left). Gene ontology enrichment bar plot of the 44 shared markers between clusters 8 and 5 (right). F. Venn diagram showing the intersect of CbO-Med211-FH cluster 6 and CbO-CHLA-01-Med cluster 4 marker genes (left). Gene ontology enrichment bar plot of the 127 shared markers between clusters 6 and 4 (right). G. Immunohistochemistry of CbO-Med211-FH co-cultures stained for GFP (green) and myogenin (red). Scalebar = 10 µm. H. Violin plot showing signature scores for the myogenic signature in different MB subgroups of a reference cohort of patient MB scRNA-Seq data^45^. I. UMAP plot of Group 3 MB reference samples in H, coloured by myogenic signature score, and labelled by their Louvain cluster. J. Barplot showing the relative proportion of myogenic (clusters 4 and 6) vs non-myogenic cells in Group 3 MB reference samples in H. K. Violin plot showing signature scores for the myogenic signature in different MB subgroups of a reference cohort of patient MB RNA-seq data^1^. L. Violin plot showing signature scores for the myogenic signature in different MB subtypes of reference cohort of patient MB RNA-seq data in K M. Survival curve showing survival data of MB patients shown in K, split by myogenic high (> median, red) or myogenic low (< median, blue) signature score.

Overall, we identify a subset of MB cells in CbO-MB exhibiting myogenic differentiation which is recapitulated in patient tumours and linked to poorer prognosis.

### CbO-MB co-cultures reveal a differential impact of medulloblastoma engraftment on cells of the tumour microenvironment

Little is known about the MB tumour microenvironment (TME), both in terms of the crosstalk between MB cells and non-neoplastic neuronal lineages and the molecular regulators of this interaction, or about the contribution of the MB-TME interplay to treatment response and recurrence in patients^69^. To investigate how MB cells affect CbO cellular complexity, we identified CbO-specific cell types in CbO-MB, annotating clusters based on label transfer using our non-co-cultured organoid as reference (Figure 7A and Figure S6A-C). The most striking difference was a depletion of mature cerebellar neurons (MatCN) which was evident in all CbO-MB co-cultures but absent in the CbO-NSC co-culture control (Figure 7B and Figure S6C,D). Selective negative impacts on neuronal populations have been shown in other brain tumour organoid co-cultures^70,71^, we therefore reasoned that MB co-culture was most likely causing neuronal death. Using IHC, we validated the presence of apoptotic neurons (cCASP3^+^ and MAP2^+^/TubIII^+^) proximal to GFP^+^ MB cells which were not present in CbO-NSC controls (Figure 7C and Figure S6E).

**Figure 7.**
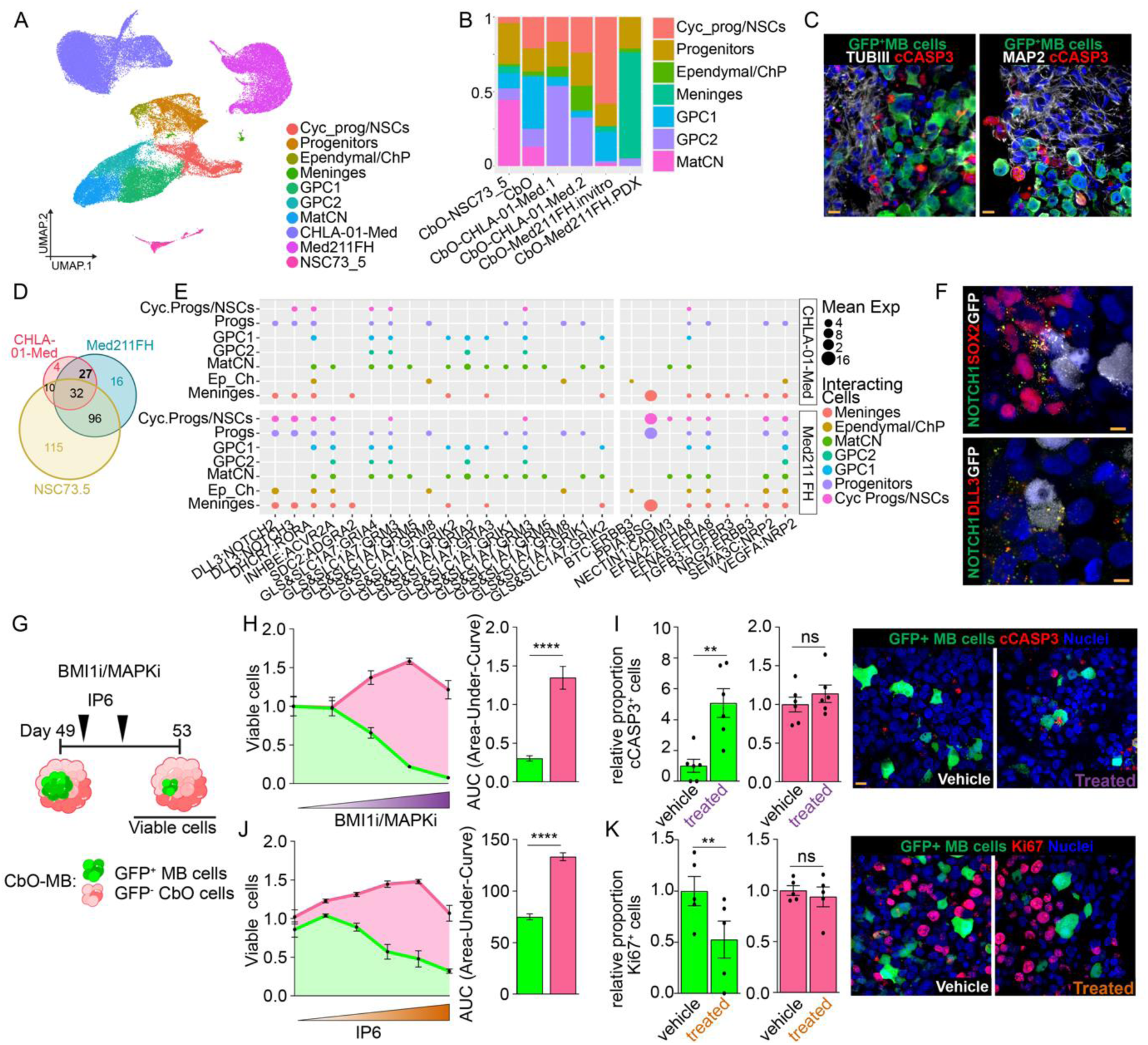
CbO-MB reveal cell-cell interactions and molecular events induced by MB growth and they represent a suitable model for drug screening. A. UMAP plot of scRNA-Seq data from D49 control CbO_61, CbO-Med211-FH, CbO-CHLA-01-Med and control CbO-NSC73_5 co-cultures, coloured by celltype. B. Barplot showing proportion of CbO cell types only present in control CbO_61, CbO-Med211-FH, CbO-CHLA-01-Med and control CbO-NSC73_5 co-cultures. C. Immunohistochemistry images of cerebellar organoids co-cultured with Med211FH for 14 days stained for GFP (green), cCASP3 (white) and MAP2(red, left) or TubIII (red, right). Scale bars = 10 µm. D. Venn diagram showing the shared significant receptor-ligand pairs modelled between co-cultured CbO-Med211-FH, CbO-CHLA-01-Med and CbO-NSC73_5 and CbO_61 cell types. Shared MB-specific receptor-ligand pairs are highlighted in bold and plotted in panel E. E. Dot plot showing significant 27 shared MB-specific receptor-ligand pairs identified by CellPhoneDB analysis interaction are separated by cell line CbO-CHLA-01-Med (top) CbO-Med211-FH (bottom), and direction MB cells as a source (left) or as a target (right). Points size corresponds to the co-expression value of the receptor-ligand pair and point colour corresponds to the interacting CbO cell type. F. Immunohistochemistry images of cerebellar organoids co-cultured with CHLA-01-Med MB cells for 14 days stained for predicted R/L interactions in E. Upper panel: NOTCH1 (green), SOX2 (red) and GFP (white); Lower panel: NOTCH1 (green), DLL3 (red) and GFP (white). Scale bars = 5 µm. G. Schematic of combination treatment with PTC209 (BMI1i) and PD329501 (MAPKi) or IP6 of CbO-CHLA-01-Med co-cultures. H. Viability assays of GFP^+^ MB cells (green) and GFP^−^ CbO cells (pink) upon 4 days of treatment with increasing concentrations of BMI1 and MAPK inhibitors. Measurement of Area Under Curve (AUC) to compare the overall response to treatment. n= 9 independent experiments. graphs report mean ± SEM, unpaired t-test ^✱✱✱✱^p<0.0001 I. Quantification (left) and representative images (right) of vehicle and treated CbO-CHLA-01 co-cultures from H stained for GFP and cCASP3. graphs report mean ± SEM, paired t-test ^✱✱^p<0.01; ns = not significant. n = 6 regions of interest quantified. Scale bars = 10 µm. J. Viability assays of GFP^+^ MB cells (green) and GFP^−^ CbO cells (pink) upon 4 days of treatment with increasing concentrations of IP6. Measurement of Area Under Curve (AUC) to compare the overall response to treatment. n= 5 independent experiments. graphs report mean ± SEM, unpaired t-test ^✱✱✱✱^p<0.0001 K. Quantification (left) and representative images (right) of vehicle and treated CbO-CHLA-01-Med co-cultures from J stained for GFP and Ki67. graphs report mean ± SEM, paired t-test ^✱✱^p<0.01; ns = not significant. n = 5 regions of interest quantified. Scale bars = 10 µm.

To characterise the molecular events mediating MB-TME crosstalk we modelled receptor-ligand (R/L) interactions in CbO co-cultures using CellphoneDB. Overall, we found that both CHLA-01-Mef and Med211-FH cells had fewer R/L interactions with CbO cells than NSC73.5 controls (Figure S6F), in keeping with MBs demonstrating limited TME reactivity compared to other brain tumours ^69^. Among the 27 cancer-specific R/L interactions shared between MB cells but absent in NSC73.5 (Figure 7D), we observed interactions mediated by the cell adhesion protein CADM3, the NOTCH signalling pathway, the receptor NRP2 as well as glutamate signalling via SLC1A7 and SLC17A7 (Figure 7E). We validated the predicted NOTCH1/DLL3 interaction between MB and cycling progenitor clusters using immunohistochemistry, revealing NOTCH1^+^SOX2^+^ CbO cells proximal to GFP^+^ MB cells, as well as co-localisation of NOTCH1 and DLL3 specifically on MB cells surface (Figure 7F).

To gain further insights into the effect of MB cells on the surrounding CNS tissue, we investigated cell type-specific transcriptional modulation caused by the presence of MB cells and identified differentially expressed genes (DEGs) (Figure S6G). GO analysis showed a downregulation of synapse and cytoskeletal pathways in GPCs whilst Ependymal/ChP cells demonstrated downregulation of cilia-related pathways and cholesterol metabolism, suggesting changes to the normal development of these populations within CbO-MB (Figure S6H). Altered neurogenesis, development and apoptosis pathways in stem/progenitor populations further highlight negative changes elicited by MB on CbO populations, suggesting a more stressed cell state. Interestingly, meningeal cells exhibited a downregulation in apoptosis and upregulation in metabolic pathways, perhaps indicating these cells better tolerate the tumour burden in CbO-MB co-cultures (Figure S6H).

Our data show that CbO-MB is a 3D model suitable for studying cell-cell interactions and molecular events/cascades induced by MB cell engraftment and identifies specific populations of CNS tissue that are more exposed to detrimental stimuli during MB formation and expansion.

### CbO-MB are an effective pre-clinical drug testing tool

Following on from the observations that CbO-MB maintain tumour cells in a molecular state closer to human samples whilst simultaneously providing insights into the functional impact of MB growth on the surrounding cerebellum, we sought to investigate whether CbO-MB could be suitable as a drug testing tool. We previously demonstrated that combination therapies extend the survival of mice xenografted with MB cells recapitulating a specific molecular signature found in a proportion of MB tumours, whereby the polycomb group gene BMI1 and the chromatin remodelling factor CHD7 are overexpressed and downregulated, respectively (BMI1^High^;CHD7^Low^).

Treatment with BMI1 (PTC-209) and MAPK (PD325901) inhibitors demonstrated a synergistic, cytotoxic effect on BMI1^High^;CHD7^Low^ MB cells whilst the phospho-inositol metabolic modulator IP6 exerted cytostatic anti-tumour efficacy^35,72^. Here, we co-cultured GFP-tagged BMI1^High^;CHD7^Low^ CHLA-01-Med cells with D35 CbO and treated CbO-MB at 14 days, a timepoint when a large population of proliferative MB cells was established (Figure 5A, B), with BMI1/MAPK inhibitors or IP6 for four days (Figure 7G). We observed a significant decrease of GFP^+^ BMI1^High^;CHD7^Low^ MB cells upon both treatments and concomitant increase in cCASP3 or decrease in Ki67 respectively, with no significant impact on CbO cells (Figure 6H-K and Figure S6I, J). This confirms that MB cells grown in 3D models recapitulate the treatment response as observed in a widely accepted pre-clinical *in vivo* model, whilst also indicating negligible side effects on CbO-MB non-neoplastic cells.

Taken together, these data show that CbO-MB are an effective 3D *in vitro* drug testing tool which perform similarly to *in vivo* xenograft models.

## Discussion

We show that CbO faithfully recapitulate the developmental stages of the human cerebellum relevant for MB initiation and contain cell lineages which display gene signature enrichment for G3 and G4 MB cells-of-origin. Genetic editing of these cells models MB onset and provides the first proof of concept that they play a role in MB pathogenesis. Moreover, co-culture of patient-derived MB cells within the CbO generates a 3D *in vitro* model which sustains MB cells proliferation and invasion while preserving their molecular identity and recapitulates the results obtained in pre-clinical *in vivo* drug testing.

Recent studies have shown the fundamental role of human-specific stem and progenitor cells in the development of G3 and G4 MB^14,15,17^, clearly exposing the limitations of mouse models and highlighting the need for new human-specific models. Putative G3 and G4 MB progenitor cells-of-origin are generated in the RL^SVZ^ upon RL compartmentalisation at 11 PCW and show a specific spatiotemporal pattern that peaks at around 14 PCW. Previously published cerebellar organoid protocols observed a correlation between two-month-old organoid cell populations and RL cell clusters from human datasets^26^ and a modest enrichment for UBC.RL^SVZ^ signature in the GNP cluster of a three-month-old organoid^14^. Here, we leveraged comparative epigenetic and transcriptomic analysis of our CbO model and human cerebellar tissue samples to accurately characterise and date the development of our model and provide the first compelling evidence that it faithfully recapitulates stages of human development known to enable MB initiation. We observed a significant enrichment for RL^SVZ^ signatures in discrete cell clusters at D35 of CbO maturation stage. Importantly, we identified specific cell clusters showing enrichment for UBC.RL^SVZ^ and PRC.RL^SVZ^ expression signatures, the latter never observed before in organoid models. TCP have been reported as putative G3/4 cells-of-origin, and we found these cells to most strongly correlate to our UBC.RL^SVZ^-high G4 MB lineage cells. Recent work suggest G3 MB originates from *PRTG*^+^;*SOX2*^+^;*MYC*^high^;*NESTIN^l^*^ow^ cells. We observed this less differentiated NSC population in our CbO, which immediately preceded the PRC.RL^SVZ^-high cluster in pseudotime/RNAvelocity trajectories. Thus, our CbO data help unify these competing theories and suggest that both putative G3 cells-of-origin are partially overlapping populations corresponding to subtly different stages along a shared G3 lineage-of-origin. Interestingly, we found a difference in distribution of EOMES^+^Ki67^+^ and PRTG^High^SOX2^+^ cells in mature CbO with respect to neural rosettes, suggesting a spatial contribution to their differing lineage commitments mimicking the events linked to neoplastic transformation of MB cells-of-origin during human cerebellar development. Therefore, our CbO model provides an unparalleled opportunity for modelling the initiation of G3 and G4 MB by providing access to their putative human-specific cells-of-origin in a 3D experimental system.

*c-*MYC amplification has been robustly demonstrated as an oncogenic driver in a subset of aggressive G3 MB, as well as reported in a proportion of G4 MB^2,73^. Previous studies have overexpressed *c-MYC* in combination with other G3 drivers *OTX2* or *GFI1* in bulk cerebellar organoids, observing an increase in proliferative cells^27^. Leveraging the in-depth developmental characterisation of our CbOs, we implemented a refined MB-modelling approach, specifically overexpressing *c-MYC* oncogene in G3/4 MB lineages-of-origin in a CbO at an MB-relevant developmental stage. This spatiotemporal inducible system generated clusters of highly proliferative cells overexpressing *c-MYC* showing an increased correlation to G3/4 MB tumours at both transcription and DNA methylation levels. Strikingly, we also observed hypomethylation of *OTX2* and *GFI1* specifically in the *pPAX6* targeted G3 lineage-of-origin model, reflecting a susceptibility of specific cells in our CbO to MB transformation as seen in patients. Genetically engineered mouse models of *MYC-* driven G3 MB derived from *in utero* electroporation of neuronal progenitors or orthotopic xenografts of cerebellar stem cells have relied on concomitant loss of *p53* function to evade cell death and induce transformation^74,75^, whilst in most human G3 MB tumours *p53* remains functional^1^. We did not observe this effect in our CbO upon *c-MYC* overexpression alone, highlighting the importance of modelling the human-specific cells-of-origin at the correct stage of cerebellar maturation. Overall, we demonstrate changes in keeping with early onset MB formation upon lineage-specific disease-defining genetic editing in human CbO, paving the way for future studies leveraging this model to dissect the events involved in the progression to full blown G3 and G4 tumours.

*In vitro* growth of MB cells isolated from patients tissue has proven difficult, particularly for G3 and G4, thus the number of reliable MB cell lines for use in 2D *in vitro* cell culture or PDX assays is limited^8,9^. Moreover, *in vitro* expansion can lead to shifts in molecular subgrouping, an issue that is particularly relevant for G3/G4 MB^4,76^. We successfully co-cultured both established and primary G3/G4 MB cell lines with CbO, showing that our 3D co-culture model represents a valuable tool to grow and maintain primary MB cell lines for which 2D culture has proven challenging by exposing tumour cells to cell-extrinsic signals from the adjacent CNS tissue. Importantly, we observed for the first time that MB cells in the 3D CbO-MB model retain an expression profile closer to that of patient tumours as compared to 2D cultures suggesting that the CbO environment is superior in supporting and preserving the molecular identity of MB cells, as has been demonstrated in co-culture models for other brain tumours^77^. Of note, despite G3/G4 MB typically manifesting post-natally, cases of congenital MB have been reported^78,79^, suggesting that the pre-natal cerebellum which we model in CbO is a permissive ecosystem for MB formation, proliferation and invasion. While multiple studies have reported the successful co-culture of primary adult glioblastoma cells with cerebral organoids^20,21^ or cerebellar organoids with established MB cell lines belonging to other subgroups^29^, this is the first time that this approach has been successfully applied to G3/4 or any primary MB cells.

Analysis of MB cells upon 3D co-culture identified clusters exhibiting neuronal and glial/stem-like phenotypes, as expected; however a cluster of MB cells displaying myogenic differentiation was also observed. Historically, this lineage of differentiation has been occasionally observed histologically in patients’ samples^67,68^, particularly in a small proportion of tumours which do not classify within the four accepted WHO subgroups^80^. Scoring of patient MBs for our CbO-MB derived myogenic signature revealed the highest enrichment in a subset of G3 patients, with myogenic differentiation being associated with poorer prognosis. Given the fidelity with which CbO-MB recapitulate this myogenic subset of G3 MB cells, they may prove a useful tool to further investigate its clinical relevance.

The characterisation of the MB TME remains limited compared to other brain tumours, with little known about the interactions or effect of MB tumours on the surrounding cerebellum^69^, however co-culture models have been successfully applied to identify TME interactions in other primary and metastatic brain tumours^81,82^. Upon MB co-culture, we observed a specific depletion of more mature neuronal CbO populations. Whether this effect is directly driven by trophic signalling of co-cultured MB cells or due to selection pressure within the CbO-MB linked to the highly proliferative nature of MB cells remains unclear. Receptor-ligand modelling predicted SLC1A7-mediated glutamate receptor signalling, which had been previously identified specifically in G3/4 MB tumours^83^. It also predicted interactions between MB and various CbO populations via the transmembrane protein NRP2 (Neuropilin-2) which had been linked to cell proliferation in an MB mouse model^84^. Proteomic studies have identified upregulation of proteins involved in TGFβ specifically in the MB TME^85^, whilst in our CbO-MB models we also saw predicted R/L interactions associated with this pathway including TGFB3/TGFBR3 and INHBE/ACVR2A^86^. We further identified and validated the interaction of NOTCH1/DLL3 between MB and CbO NSC populations, previously described to be involved in MB’s prognosis, tumour microenvironment and metastasis^87,88^. Overall, we demonstrate our CbO-MB co-culture is a useful model to dissect potential TME mediators linked to MB progression and metastasis.

Finally, our CbO-MB model recapitulates the therapeutic response upon treatment with BMI1/MAPK inhibitors and IP6 as previously demonstrated in xenograft models, hence supporting its potential value as a system for testing MB subgroup specific and signature-matched therapeutic approaches. Notably, CbO-MB represent a faster, scalable and more cost-effective model system as compared to expansion of MB cells in immunodeficient mice, whilst also offering the possibility to evaluate toxicity on cerebellar cells induced by anti-tumour therapy, hence could be used in precision medicine modelling to identify new treatment options that are still lacking for G3/G4 patients.

In conclusion, we have established a new human cerebellar organoid model which enabled access to human G3/G4 MB cells-of-origin and provided proof of concept that genetic editing of these cell lineages leads to changes defining MB onset. Moreover, CbO sustained G3/G4 MB cell growth in a co-culture system, enabling the identification of a myogenic signature linked to poor prognosis as well as modelling in 3D the tumour microenvironment. Finally, we show its potential value as a novel scalable pre-clinical screening platform for precision medicine.

## Material and Methods

### Cell Culture

Full characterisation of the two EPSC lines used in this study has been reported previously^21^. Briefly, EPSC19 and EPSC61 were derived from fibroblasts isolated from the healthy dura mater of two female patients harbouring no somatic mutations and reprogrammed based on established protocols^30^. EPSCs were cultured as described previously^21^ on plates coated in Geltrex (Life Sciences) in mTEsR plus basal media (StemCell Technologies). During routine passaging, cells were detached with 0.1 mM EDTA (Life Sciences) and re-plated with media supplemented with 10 μM Y-27632 Dihydrochloride (PeproTech, #1293823). CHLA-01-Med and CHLA-01R-Med MB cell lines purchased from ATCC, were cultured in suspension in Gibco DMEM-F12 media (Life Technologies) supplemented with 20 ng/mL human recombinant FGF and 20 ng/mL human recombinant EGF (Peprotech), 2% B-27 supplement (Invitrogen) and 1% Penicillin-Streptomycin. ICb1299 MB cells obtained from Dr Xiao-Nan Li, Baylor College of Medicine, Texas Children Cancer Centre, USA^11,31^ were cultured semi-adherently in Gibco DMEM media (Life Technologies) supplemented with 10% foetal bovine serum (Gibco) and 1% Penicillin-Streptomycin (Life Sciences). Med211-FH cells tagged with GFP were cultured in Neurobasal Plus media (Life Sciences) supplemented with N2 (Bio-techne), B27 (Bio-techne), 75 µg/mL BSA, 2 µg/mL heparin (StemCell Technologies), 10 ng/mL human recombinant FGF and 10 ng/mL human recombinant EGF (Peprotech).

All cell lines were confirmed mycoplasma negative prior to experimental procedures and cultured in humidified incubators at 37°C with 5% CO2 until ∼80% confluent before passaging.

### Cerebellar Organoid Culture

Cerebellar organoids were cultured based on a previously published protocol^22,23^. Briefly, EPSCs at 80% confluency were detached using gentle cell dissociation reagent (StemCell) and plated into 96-well ultra-low attachment V-bottomed plates (PHCBI) in mTEsR plus basal media (StemCell) supplemented with 10uM Y-27632 Dihydrochloride (PeproTech, #1293823) at 6,000 cells per well. On Day 1, media was replaced with mTEsR plus basal media. On Day 2, media was replaced with growth-factor free chemically defined media (gfCDM) containing: IMDM (Life Technologies)/Ham’s F-12 (Life Technologies) 1:1, 1% v/v chemically defined lipid concentrate (Life Technologies), 450 μM monothioglycerol (Sigma), 15 μg/ml apo-transferrin (Sigma), 5 mg/ml crystallization-purified BSA (Sigma), 50 U/ml penicillin/50 μg/ml streptomycin and 7 μg/ml insulin (Sigma). On Day 2-7, 50 ng/ml human recombinant FGF2 (PeproTech, #100-18B) and 10 μM SB431542 inhibitor (Life Sciences, #S4317) were added to gfCDM. On Day 7, organoids were transferred to 6-well ultra-low attachment plates (Corning) on an orbital shaker (90 rpm, Thermo Scientific, #88881102) in a humidified incubator at 37°C with 5% CO2 where they were maintained for the remainder of the protocol. For Day 7-14, 33.3 ng/ml human recombinant FGF2 and 6.67 μM SB431542 inhibitor were added to gfCDM, then 100 ng/ml human recombinant FGF19 (PeproTech, #100-32) was added to gfCDM for Day 14-21 of culture. From Day 21, organoids were cultured in complete neurobasal media containing: Neurobasal medium (Life Technologies), GlutaMax I (Life Technologies), N2 supplement (Life Technologies), and 50 U/ml penicillin/50 μg/ml streptomycin. 300 ng/ml human recombinant SDF1 (PeproTech, #300-28A) was added to complete neurobasal media from Day 28-35.

### CbO editing strategy

To induce *c-MYC* expression in putative MB lineages-of-origin within developing CbOs, three lentivirus constructs were designed utilising the doxycycline inducible Tet-On system^32^. To target specific CbO lineages, the *rtTA* sequence element and *BFP* tag were placed under the control of the *EOMES* promoter (p*EOMES*) or a *PAX6* mini promoter (Ple260^33^) designed for CNS expression (p*PAX6*). In addition, each construct contained the puromycin resistance gene under the control of a constitutively expressed promoter (*mPGK)*, giving *pPAX6-rtTA-BFP_mPGK-PuroR* or *pEOMES-rtTA-BFP_mPGK-PuroR*. The third construct contained the *c-MYC* sequence and *GFP* tag under the control of the tetracycline responsive element (*TRE)* as well as blasticidin resistance gene under the control of *mPGK,* giving *TRE-c-MYC-GFP_mPGK-BlastR.* Lentiviral constructs were produced as described previously^34^ and titrated using Lenti-X GoStix Plus as per manufacturer instructions (Takara Bio). EPSC19 were infected with *pPAX6-rtTA-BFP_mPGK-PuroR* or *pEOMES-rtTA-BFP_mPGK-PuroR* (MOI = 1), negative selection for 24 hours performed with puromycin (1.25 µg/mL) then co-infected with *TRE-c-MYC-GFP_mPGK-BlastR* construct (MOI = 1) and Blasticidin (5 µg/mL) negative selection performed for 48 hours. Cerebellar organoid protocol was then begun as detailed above. From day 35, doxycycline (2 µg/mL) was added to culture media and CbOs monitored for 4 weeks.

Two sets of controls were included, non-induced CbOs harbouring the full editing system (no doxycycline control) as well as CbOs derived from EPSC infected with the *pPAX6-rtTA-BFP_mPGK-PuroR* or *pEOMES-rtTA-BFP_mPGK-PuroR* constructs only and treated with doxycycline from day 35 onwards (doxycycline+ control). On day 63, edited CbO were taken for scRNA-seq analysis, IHC (both detailed below) or dissociated, FACS sorted for BFP or GFP using BD FACSAria III Cell Sorter then DNA extracted and submitted for EPICv2 array (outlined below).

### CbO-MB co-culture

Prior to co-culture, G3/4 MB cell lines were infected with shCHD7 lentivirus containing GFP as previously published^35^. On Day 35 of CbO culture, organoids were placed in 96-well ultra-low attachment U-bottomed plates (Corning) in a 1:1 mix of complete MB cell media and complete neurobasal media with 100,000 MB cells. For Med-211-FH cells previously cultured as PDX only, cells were thawed then underwent mouse cell depletion using Miltenyi mouse cell depletion kit (#130-104-694) according to manufacturer instructions then seeded onto CbO as described above. The following day, half media from each well was replaced with complete neurobasal media. On Day 37, the entire contents of each well was transferred to a 24-well ultra-low attachment plate (Corning) with 1ml complete neurobasal media. On Day 4, total media was replaced with complete neurobasal media to remove non-infiltrating cells and organoids placed back on the orbital shaker (90 rpm) in a humidified incubator at 37°C with 5% CO2. Co-cultures were maintained for 14 days in total from MB cell seeding before taking for drug assays or scRNA-Seq.

### Human Cerebellar FFPE Samples Processing

12 human cases from Great Ormond Street Hospital (GOSH) at pre-natal developmental timepoints (19-37 post-conception weeks) were selected as having normal cerebellar histology and consent for research. Areas of formalin-fixed paraffin-embedded (FFPE) tissue containing cerebellar material were identified and manually macro dissected according to the marking shown in Figure S2A. Following dissection, tissue was kept at −20°C until downstream processing. In addition, 11 samples of fresh frozen cerebellar tissue from earlier developmental timepoints (Carnegie stage 22-17 post-conception weeks) were obtained from the Human Developmental Biology Resource (HDBR).

### DNA/RNA Extraction

For cell culture samples (CbO, EPSC and Med211-FH), DNA and RNA were extracted using the RNA/DNA/Protein Purification Plus kit (Norgen, #47700) following manufacturer protocol. For FFPE human cerebellar samples, initial deparaffinization and DNA extraction was performed using FFPE RNA/DNA Purification Plus Kit (Norgen, #54300). Samples were incubated with Proteinase K overnight at 37°C to increase yield, after which DNA extraction was performed as per manufacturer instructions. For fresh frozen human samples, extraction was performed using RNA/DNA Purification Plus Kit (Norgen, #47700) as per manufacturer instructions.

### RT-qPCR

All cDNA synthesis was performed using the SuperScript III Reverse Transcriptase kit (Invitrogen) according to the manufacturer’s protocol. Analysis of gene expression was carried out using the Applied Biosystems 7500 Real-Time PCR with SYBR Green PCR MasterMix (Applied Biosystems) according to standard protocols. All qPCRs were performed in triplicate with 10ng cDNA. Average Ct values for target genes were normalised to average Ct of ACTIN B and GAPDH housekeeping genes before calculating fold change. Primers used are listed in Table 1.

**Table 1.**
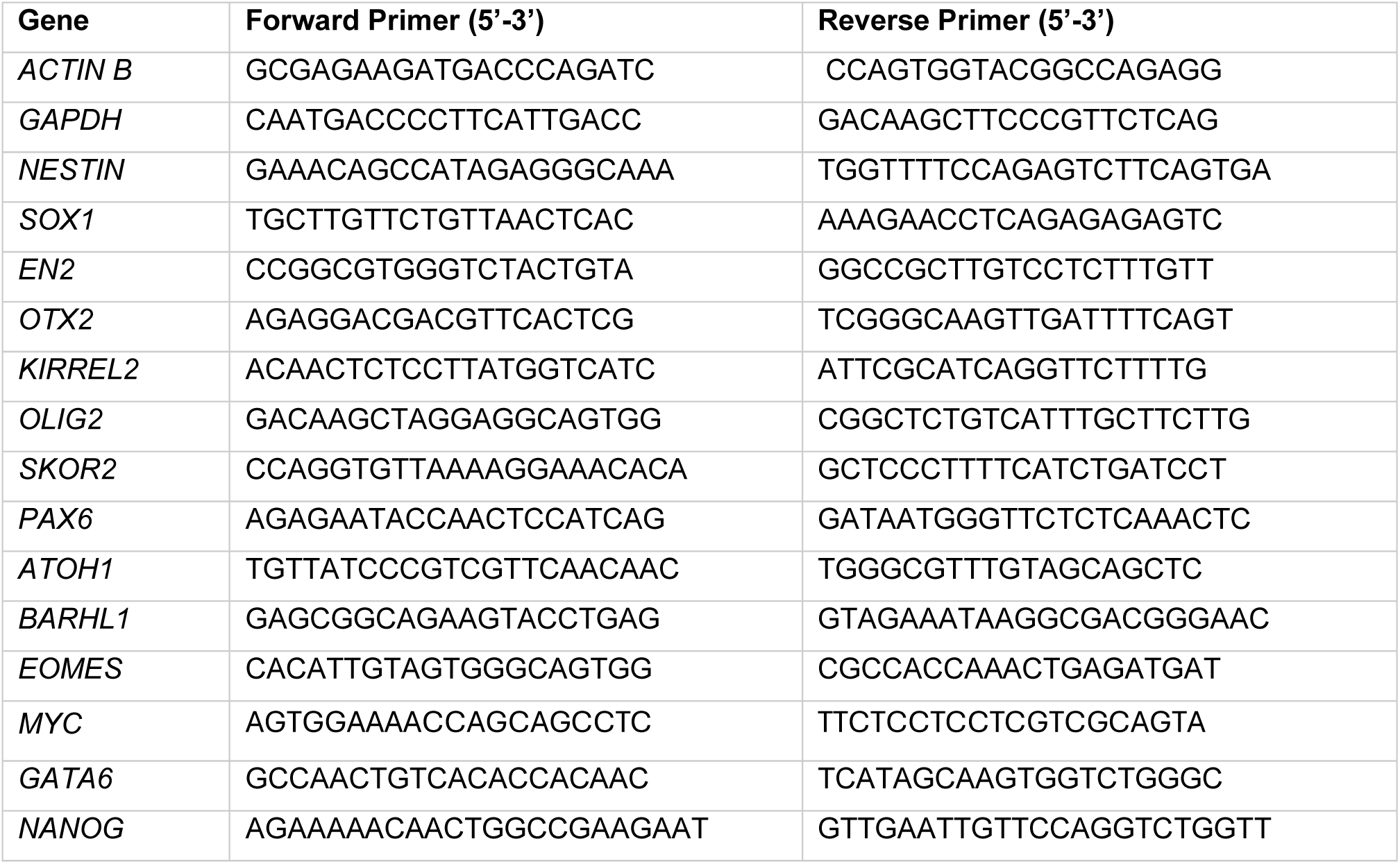
Primers used for RT-qPCR reactions.

### Immunohistochemistry (IHC)

Organoids were collected and fixed in 4% PFA in PBS, dehydrated in 30% sucrose in PBS and embedded in blocks of 7.5% gelatin, 10% sucrose in PBS. Blocks were then snap frozen in isopentane at −55°C before cryosectioning at 15 μm sections. Prior to staining, sections were de-gelatinised in warm water before permeabilisation in 4% normal donkey serum with 0.25% Triton-X in PBS for 2 hours. Sections were then washed in PBS before incubation with primary antibodies at 4°C overnight diluted in 4% normal donkey serum in PBS. A list of primary antibodies and dilutions are shown in Table 2. Sections were then washed 3 times in PBS followed by incubation for 3 hours at room temperature with corresponding AlexaFluor™ secondary antibodies at 1:500 dilution in PBS. After a final 3 PBS washes, sections were mounted with coverslips using ProLong™ Gold antifade reagent with DAPI (Thermo Fisher Scientific, P36931). Images were taken using Zeiss 800 Laser Scanning Confocal microscope with Zeiss ZEN software v2.3.

**Table 2.** Primary antibodies and dilutions used for immunohistochemistry.

| Primary Antibody | Dilution |
| --- | --- |
| SOX2 (abcam, ab97595) | 1:500 |
| OLIG2 (Santa Cruz, sc-19967) | 1:200 |
| CALB (Swant, CB300) | 1:200 |
| NeuN (Chemicon, MAB377) | 1:500 |
| EOMES (Chemicon, AB15894) | 1:200 |
| KI67 (abcam, ab15580) | 1:500 |
| GFP (Chemicon, AB16901) | 1:400 |
| cCASP3 (Cell Signalling, 9664) | 1:500 |
| MAP2 (Sigma, M4403) | 1:200 |
| TubIII (abcam, ab7751) | 1:200 |
| Myogenin (abcam, ab1835) | 1:100 |

### CbO-MB drug assays

On Day 0 of drug assays, CbO-MB were transferred to new 24-well ultra-low attachment plates (Corning) containing 500 μL complete neurobasal media plus drug (PTC-209/PD325901 inhibitors or IP6) or vehicle (DMSO or PBS respectively) then placed back on orbital shaker where they remained for the duration of the assay. A drug and media top up was performed on day 2.

On day 4, the entire content of each well was collected and the Neural Dissociation kit (MACS Miltenyi Biotec) was used to dissociate treated CbO-MB co-cultures into single cell suspension. Samples were then stained with the Zombie NIR™ Fixable Viability Kit (Biolegend) at 1:300 dilution, after which all samples were analysed using the BD LSRII flow cytometer. GFP-expressing cells were detected with the blue excitation laser at 488nm, while the Zombie NIR positive cells were detected with the red excitation at 633nm. Upon gating of single cells, Zombie NIR positive cells were gated as dead, with GFP fluorescence distinguishing viable MB from CbO cells. All subsequent FACS analysis was performed in FlowJo v10.6.2.

### Dissociation of CbO and CbO-MB for scRNA-Seq

Pooled samples of CbO were taken at day 7 and day 35, as well as edited CbO at day 63 and CbO-MB at day 49. First, organoids were dissociated using the Neural dissociation kit (MACS Miltenyi Biotec) into single cell suspension. Subsequent debris removal was then performed using Debris removal solution according to manufacturer protocol (Miltenyi Biotec, #130-109-398) to obtain a clean sample of highly viable cells. Cells were resuspended in loading buffer (0.04% BSA in PBS), strained through a 70 μm cell strainer and counted using a hemacytometer with trypan blue staining. At least 10,000 cells per sample were taken for processing using 10X Genomics kits (detailed below).

### RNA sequencing

Med211-FH RNA from cells cultured *in vitro* was sequenced at Novogene with paired-end, 150bp sequence reads. Library preparation (mRNA, poly-A enrichment) and sequencing was performed on the NovaSeq6000 platform. Reads were quality trimmed using trimgalore v0.6.5 and aligned to reference genome GRCh38.release109 using STAR v2.7.9a^36^. Reads were counted using featurecounts function and annotated with ENSEMBL Homo_sapiens.GRCh38.109.gtf.

### scRNA-Seq Analysis

Library preparation was performed at the Single Cell Genomics Facility (UCL, Cancer Institute) using the Chromium Next GEM Single Cell 3’ Kit v3.1 (10X Genomics, 1000268) along with the Chromium Next GEM Chip G Single Cell Kit (10X Genomics, 1000120) according to the manufacturer’s instructions. Pooled samples libraries were sequenced with lane sequencing on Novaseq 6000 S4 (Novogene). The Cell Ranger 2.0.1 pipeline was used to align reads to the GRCh38 human reference genome and produce count matrices for downstream pre-processing and analysis using the Seurat v5.0.1 R package^37^.

### UMAP calculations and cell type annotations

For analysis of CbO samples we implemented stringent quality filtration, only cells with fewer than 5×10^4^ total reads, lower than 10% mitochondrial reads, more than 750 and less than 6500 detected features were retained for analysis. Expression values were library size corrected to 10,000 reads per cell and log1p transformed, with Principal component analysis (PCA) performed on the scaled data for the top 2000 variable genes. Batch correction between cell lines was performed on principal components using Harmony^38^. Uniform Manifold Approximation Projection (UMAP) embeddings, Nearest Neighbours and cell clusters were then calculated in harmony-corrected PCA space using 35 dimensions, and cells were clustered using FindClusters(). For Day 7 COs, FindClusters() resolution was set at 0.2, and for Day 35-49, where the UMAP indicated a greater degree of cell type complexity, we instead set a resolution of 0.5 and used FindSubCluster(resolution = 0.1, algorithm = 1) to separate cluster 7 into two distinct subclusters 7_0 and 7_1. Cluster marker genes were calculated using a Seurat’s in-bult Wilcoxon Rank Sum test (logfc.threshold = 0.25, min.pct = 0.1, only.pos = T), and differential expression analysis across experimental conditions was performed using MAST^39^. Initial clustering identified two clusters 13 and 14 which represented hypoxic/glycolytic cells, common to all human brain organoids^40^. As has been done previously^41^, these cells were excluded from analyses and a new UMAP was calculated.

For analysis of engineered CbO and CbO co-cultures containing CHLA-01-Med and Med211FH MB medulloblastoma (MB) cells or control NSC37.5 cells, we changed the detected feature threshold greater than 500 and less than 7500, with no batch correction was necessary. For engineered CbO UMAPs and Nearest neighbours were calculated using 30 PCA dimensions and FindClusters() was run with a resolution of 0.5. For CbO co-cultures we used 15 PCA dimensions and a resolution of 0.35. Malignant CHLA-01-Med and Med211FH MB cells were distinguished from co-cultured CbO cells based on Leiden clustering and inferred copy number variants, with co-cultured NSC73.5 identified based on clustering and marker gene expression alone. InferCNV was run on CbO-CHLA-01-Med and CbO-Med211FH samples using D49 CbO cells as a reference, the “subclusters” analysis mode, and setting the following parameters: cutoff=0.1, k_obs_groups = 3, leiden_resolution = 1×10^−5^. For annotating engineered CbO and organoid cells from CbO-MB co-cultures we used the processed R object from the D35-49 CbO (Fig1B) as a reference, and the default settings of Seurat FindTransferAnchors(), TransferData() and AddMetaData() functions. We then annotated new cluster names based on their composition of transferred cell type labels Fig S2D-G.

Finally, for the analysis of MB cells alone, cells CHLA-01-Med and Med211FH identified by InferCNV were subset, and reanalysed using the standard Seurat pipeline; UMAPs and nearest neighbours were calculated using 20 PCA dimensions and FindClusters() was run with resolution=0.225.

### scRNA-Seq Analysis: Comparison to reference datasets

To approximate the transcriptomic age of our CbO we compared data from this study, and three other cerebellar organoid studies: Nayler et al^25^, Chen et al^26^, and Atamian et al^24^ against four datasets derived from developing foetal cerebellum: Aldinger et al^42^, Luo et al^16^, Zhong et al^43^, and Sepp et al^44^. First, pseudo-bulk transcriptomic data were generated from aggregated single cell counts for all samples, and expressed as log(counts per million +1). Data was merged and batch correction between different sequencing methodologies (scRNAseq vs snRNAseq) was performed using Combat() from the sva.3.48 package. We next derived an ‘ageing-signature’ as the top 200 genes correlated with age across the foetal cerebellum samples. This signature was then used to compute the average Pearson correlation between CbO and foetal cerebellum samples at different weeks of development.

To aid with D35/49 CbO cell type annotations, we compared cluster marker signatures against those from the four foetal cerebellum datasets. In each case we obtained published count matrices, UMAP-embeddings and author-defined cluster meta data, and used them to perform a Wilcoxon Rank Sum test with Seurat’s FindAllMarkers() command (logfc.threshold = 0.25, min.pct = 0.1, only.pos = T) to obtain the list of marker genes for their cell types. Then, for each dataset and marker gene, cluster specificity scores were computed (mean normalized counts per cluster/total mean normalized counts) – with overlapping gene signature specificity scores compared across studies by Pearson correlation. We performed the same analysis on CHLA-01-Med and Med211FH cells to determine each MB cluster’s closest analogues from developing foetal cerebellum samples.

To compare CHLA-01-Med and Med211FH MB cells in co-cultures to patient tumours, we generated pseudo-bulk transcriptomic data from aggregated single cells counts from our co-cultured MB samples, and 36 patients in Hovestadt et al^45^. Combat batch correction was performed between log(cpm+1) transformed data, and Pearson correlation between samples were calculated across the top 2500 most variable genes (ranked by standard deviation). Pearson correlation across the same gene list was also used to query the relative similarity between patient tumours and CbO co-cultured CHLA-01-Med cells profile here, and CHLA-01-Med cells we previously cultured *in vitro* and profiled by RNAseq^35^. For Med211FH MB cells, we compared CbO co-cultured to RNAseq data from an *in vitro* sample drawn from the literature^17^ and a second *in vitro* sample profiled as part of this study.

### scRNA-Seq Analysis: Reference gene set/signature scoring

For reference signature scoring, average gene module expression was calculated for each single cell, subtracted by the aggregated expression of a random control set of features selected from the same average expression bins as the query genes^46^ using Seurat’s AddModuleScore() function. G/S and G2M gene modules are included in the Seurat v5.0.1 package; G3 RL.PRC and G4 RL.UBC Medulloblastoma cell of origin signatures were taken from Smith et al^14^; Group 3 and 4 MB tumour signatures were taken from Vladoiu et al^47^; WNT, SHH and G3/4 MB cell state signatures were taken from Hovestad et al^45^. For gene set enrichment analysis, gene sets were obtained from MolSigDB^48^ and computations were performed using the R package fGSEA v4.4^49^. Gene ontology enrichment analysis of was performed on upregulated (p.adj<0.05, and avg_log2FC>2) genes in the MYC^oe^ cluster of engineered CbOs using the topGO v2.52.0 and org.Hs.eg.db v3.17.0 R packages.

Processed cell count and metadata from patient-derived MB samples was accessed from Hovestadt et al^45^ and reanalysed in Seurat using default settings. The resulting R object was then used as a reference, with the default settings of Seurat FindTransferAnchors(), TransferData() and AddMetaData() functions to assign MB Group3 and Group4 prediction scores to the co-cultured CHLA-01-Med and Med211FH MB cells. To generate a Myogenic MB state signature, we took the intersect of significant MB marker genes with avg_log2FC>1 from cluster 4 (CHLA-01-Med) and cluster 6 (Med211FH). The processed reference R object from Hovestadt et al^45^ was then scored for the Myogenic MB state signature as detailed above using the the AddModuleScore() function. Next, the Hovestadt et al^45^ data was subset to group 3 samples, before Harmony^38^ batch correction between samples was performed. New UMAP embeddings, Nearest Neighbours were then calculated in harmony-corrected PCA space using 15 dimensions, and clusters were identified using a resolution of 0.5. We also inspected a large reference dataset of 763 bulk MB RNAseq samples (Cavelli et al^1^) for expression of the Myogenic MB state signature using the GSVA 1.48.3 R package^50^. Survival analysis was performed on patients with complete data (i.e. that had died) using the survival_3.8-3 and survminer_0.5.0 R packages and splitting patients into the Myogenic Signature High or Low groups based on the median signature score.

### Cell lineage trajectory analysis

For RNAvelocity analysis, velocyto^51^ was used to calculate spliced and unspliced transcriptomes, before subsequent analysis using scVelo^52^, on Seurat-defined cell types and UMAP embeddings, using default parameters. For single cell lineage trajectory analysis, we imported the processed Seurat object into the Monocle3 R package. The ‘learn_graph’ function was applied to infer single cell trajectories setting close_loop = TRUE, minimal_branch_len to 10 and euclidean_distance_ratio to 1.5. The ‘order_cells’ function was then used to set Progenitors and Cyc_Progenitors as the root of states of pseudotime. Next we subset data to cell types involved in the G3 trajectory (early_RL/NSCs, glut_progenitors and matCN2) or G4 trajectory (NSCs, GN/UBC_progs and GN/UBC)^53^. For each subset, the ‘learn_graph’ function was applied again to infer single cell trajectories setting minimal_branch_len to 10 and the ‘order_cells’ function was used to select early_RL/NSCs and NSCs as the root states of pseudotime. Differential gene expression was calculated across single cell trajectories using Moran’s I test. For each trajectory, the top 2500 most differentially expressed genes over pseudotime were selected, and smoothened, alra-imputed^54^ expression over pseudotime was plotted.

### Receptor Ligand Modelling

For receptor-ligand analysis, cell type meta data and count data for the top 15000 most highly variable genes in the CbO-MB dataset was input into CellphoneDB v5^55^ and run using the statistical_analysis mode.

### DNA methylation analysis

All CbO and human cerebellar DNA samples underwent standard BSC conversion at UCL Genomics facility (UCL, Great Ormond Street Institute of Child Health) and were submitted for Illumina Methylation EPIC v2 array following standard protocols. Quality control was performed and probe beta values were extracted from raw .idat files using wateRmelon R package^56^ under reference genome hg38. Probes with detection p-values greater than 0.05 were removed using pfilter(), as well as probes associated with sex chromosomes, mitochondrial chromosomes and single nucleotide polymorphisms. Adjusted Dasen normalisation was then performed.

### Cerebellar Organoid developmental staging (DNA methylation)

To stage the development of CbO compared with human cerebellar samples, an ‘ageing signature’ approach was applied, with the top 1% of probes whose beta values best correlated to age in human cerebellar samples selected for subsequent correlation with CbO probe betas. Combat batch correction was first performed between CbO and human samples before top 1% probe selection and correlation to CbO. To corroborate this approach, CbO methylation data adapted to EPIC v1 array probe annotation was also passed to the foetal brain clock (FBC), an R package specifically trained on foetal brain samples during development to estimate their epigenetic age^57^.

### Differentially and variably methylated probes and regions analysis

Differentially methylated probes (DMPs) and regions (DMRs) were calculated pairwise for CbO samples between comparison groups using the R package DMRcate^58^ with FDR threshold of 0.01. Additional criteria for DMR identification were set such that DMRs must contain at least 6 CpG sites with <400 base pair separation. Variable methylated probes (VMPs) and regions (VMRs) across CbO and human development timepoints were identified within the DMRcate package using the same thresholds as stated above for DMR analysis and the top 5% VMP/VMRs defined as significant. VMRs were then annotated to genes (VMGs), after which significant VMGs were clustered based on z-scores of average beta-values of the probes contained within them and taken for pathway analysis.

### DNA methylation: Comparison to reference datasets

For human rhombic lip (RL) and external granular layer (EGL) correlation, raw .idat files for micro-dissected RL and EGL samples were accessed from GSE207266^14^ and processed using the same raw data processing pipeline adapted for EPIC v1 array annotation before downstream analysis. DMPs were identified between each cerebellar compartment (n=7) as stated above with FDR threshold of 0.01. DMPs were then subset to CbO average probe betas adapted to EPIC v1 array annotation and correlated to each cerebellar compartment.

For CbO editing samples, reference 450K methylation data of 763 MB samples were accessed from GSE85218^1^. CbO datasets were trimmed to probes present in both 450K and EPICv2 array and clustering of CbO and MB samples was performed based on a published probe set shown to distinguish between MB subgroups^59^ (34/48 CpG probes present in QC filtered datasets). DMPs were calculated as described above on the trimmed CbO dataset then correlated to patient MB samples.

### scRNA-Seq/DNA methylation integration and pathway analysis

Differentially expressed genes (DEGs) between Day 7 and Day 35 CbO scRNA-Seq data with adjusted p-value < 0.05 and absolute log2FC > 0.25 were integrated with genes mapping to DMRs from methylation data at matched timepoints with p-value < 0.05 and greater than a 10% imposed threshold in methylation difference. Genes demonstrating concordant hypermethylation and down-regulation or hypomethylation and up-regulation were taken for subsequent pathway analysis. For pathway analysis, gene lists were input into Gprofiler ^60^ and pathways calculated based on a Benjamini-Hochberg FDR < 0.05. Pathway visualisation was performed using CytoScape ^61^. For scRNA-Seq upon MB co-culture pathway analysis, UMAP clusters were grouped based on cell developmental state (un/differentiated) and effect of MB co-culture on cluster percentage (increase/decrease). Common pathways from each group were then identified and the top 100 most significant (adjusted-p<0.05) Reactome and KEGG pathways were plotted. For VMR pathway analysis, the top 100 most significant (adjusted-p<0.05) GO:Biological Process and GO:Molecular Function pathways were plotted for human and CbO VMG clusters, whilst for overlapping pathways with concordant DMR/DEG integration of Day 7 vs Day 35 CbO samples, GO:Cell Compartments were also plotted.

## Ethics

Ethical approval for the work on human cerebellar samples was obtained from Brain UK, reference number 22/010.

## Acknowledgements

This work is funded by grants from Brain Tumour Research (Centre of Excellence award to S.M.), Cancer Research UK (C23985/A29199 programme award to S.M.), Barts Charity (MGU0447 programme grant to S.M.). We thank Gary Warnes (Blizard Flow Cytometry Core Facility), Maeve McLaughlin (Blizard Advanced Light Microscopy Facility), Imran Uddin (Single Cell Genomics Facility, UCL, Cancer Institute), Myrianni Constantinou and Sara Lucchini for sharing their expertise. The collection and annotation of FFPE human tissue samples are funded by the NIHR GOSH BRC. The collection of fresh frozen human tissue samples is funded by the MRC-Wellcome Trust HDBR.

## Author contribution

SB and SM conceived and supervised the project; TW, SB and SM designed the experiments; TW performed, analysed and interpreted all experiments and data; SB analysed and interpreted all experiments and data; JN analysed and interpreted scRNA-seq data; TW and YX analysed DNA methylation data; OO, TM and AM identified and provided suitable human foetal cerebellar samples; BD and LD shared G3 MB primary cells, datasets and expertise; NRZ contributed expertise for DNA methylation analysis; SM secured financial support; TW, SB, JN and SM wrote the paper with contributions from all authors.

## Data Availability

The authors declare that all the data supporting the findings of this study are available within the article and its supplementary files. The scRNA-seq and DNA methylation datasets generated in this study and processed data are available in the NCBI Gene Expression Omnibus database (GSE263652 and GSE263653 respectively) or are available from the corresponding authors upon reasonable request. Publicly available datasets used in the study were retrieved from relevant databases as outlined in materials and methods.

## Code Availability

The authors declare that no custom code was used in this study. A full description of analyses including functions and packages used are stated in the materials and methods. Specific code will be made available from the corresponding authors upon reasonable request.

## Supplementary Figures and Legends

**Supplementary Figure 1.**
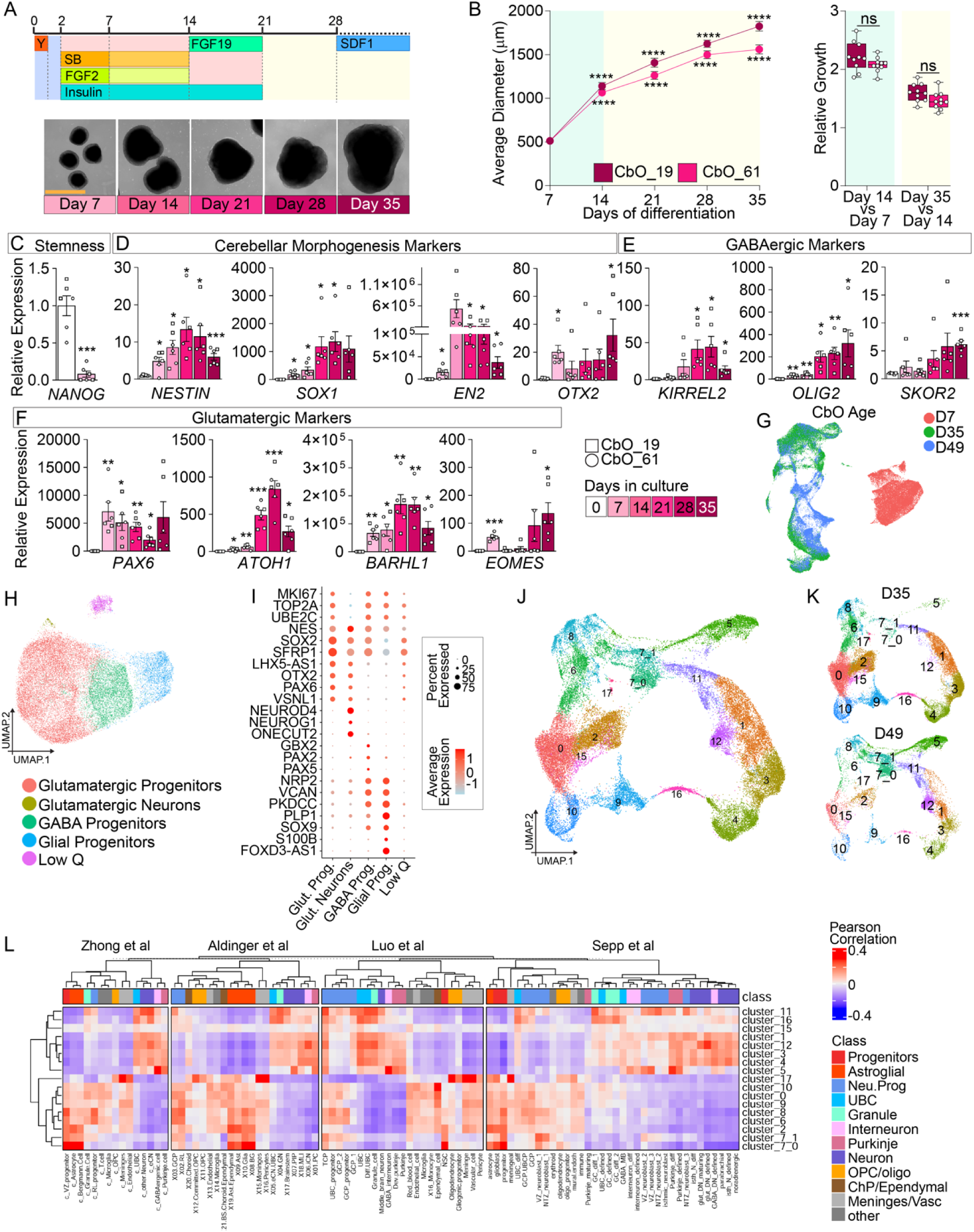
CbO displays sustained growth and maturation. A. Schematic of protocol used to generate and differentiate CbO, (see Methods, Y: Y-27632 Dihydrochloride and SB: SB431542) and representative brightfield pictures of CbO_61 upon 7, 14, 21, 28 and 35 days of differentiation. Scale bar = 1000 µm. B. Diameter analysis of CbO obtained from two independent EPSC lines (CbO_19 and CbO_61) across 35 days of maturation (left) and analysis of relative growth comparing Day 14 to Day 7 or Day 35. Asterisks indicate p-values between given time point and Day 7, two-way ANOVA, ^✱✱✱✱^p<0.0001, ns: not significant. n = 10 independent CbO average diameters per timepoint. C. qPCR analysis showing quantification of *NANOG* expression in CbO_19 and CbO_61 across 35 days of maturation. D. qPCR analysis showing quantification of *NESTIN, SOX1, EN2* and *OTX2* expression in CbO_19 and CbO_61 across 35 days of maturation. E. qPCR analysis showing quantification of *KIRREL2* and *SKOR2* expression in CbO_19 and CbO_61 across 35 days of maturation. F. qPCR analysis showing quantification of *PAX6, ATOH1, BARHL1* and *EOMES* expression in CbO_19 and CbO_61 across 35 days of maturation. Relative RNA level was calculated as 2^(-dCT) values normalised to EPSC baseline (Day 0). All graphs report mean ± SEM, Welch’s unpaired t-test, ^✱^p<0.05, ^✱✱^p<0.01, ^✱✱✱^p<0.001. non-significant comparisons are not shown. Data points show average values for n = 3 independent batches of CbO per timepoint. G. UMAP plot of scRNA-Seq data from all CbO samples submitted at Day 7 (CbO_19 and CbO_61), 35 (CbO_19 and CbO_61)and 49 (CbO_61) of differentiation coloured by timepoint. H. UMAP plot of scRNA-Seq data from CbO at Day 7 of differentiation coloured by annotated Louvain clusters. I. Dot plot showing key marker genes used to identify major cell types in CbO at Day 7 of differentiation. J. UMAP plot of scRNA-Seq data from CbO samples at Day 35/49 of differentiation, identifying 17 clusters and sub clustering of cluster 7 into 7_0 and 7_1. Low quality hypoxic clusters 13 and 14 not shown. K. UMAP plot of scRNA-Seq data shown in K, split by Day 35 (upper) and Day 49 (lower) timepoints. L. Heatmap showing correlation of CbO D35/49 clusters identified in K to clusters annotated by cell type of four independent scRNA-seq datasets of the human developing foetal cerebellum^1–4^.

**Supplementary Figure 2.**
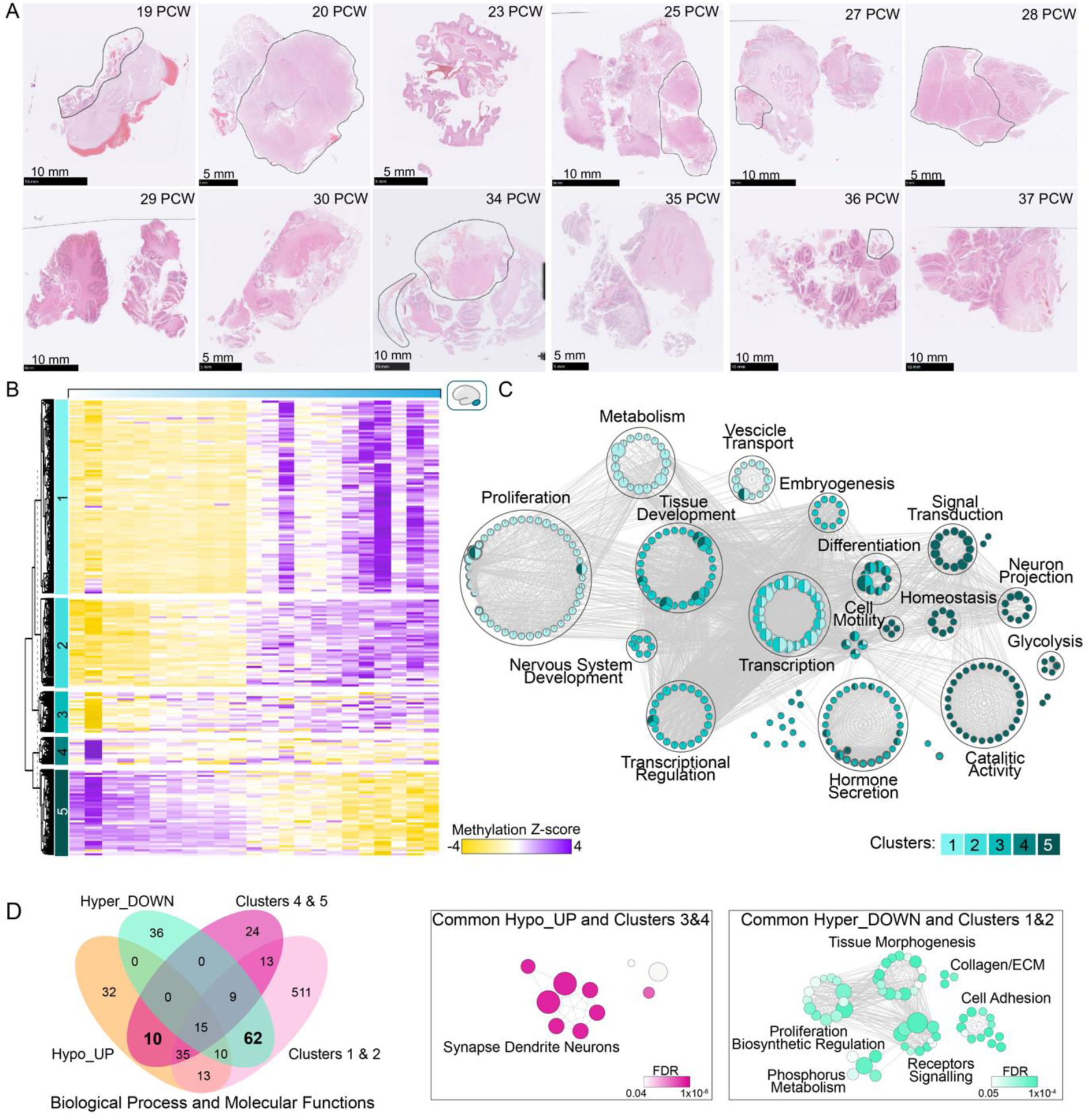
Staging development of CbO comparatively to human cerebellum reveals conserved epigenetically regulated genes and pathways. A. H&E staining of FFPE samples from human foetal cerebellum. Non-cerebellar tissue excluded from the microdissection is circled. Developmental stage in post-conception weeks (PCW) and scale bars are shown. B. Heatmap showing k-means clustering (n=5) of significant variable methylated genes across human foetal cerebellar development from Carnegie stage (cs) 22 to 37 post-conception weeks (pcw). Z-scores for average beta values of probes contained within each gene are plotted. C. Bubble plot showing top 100 significant (adjusted p-value < 0.05) GO Biological Processes and Molecular Functions of genes from 5 clusters shown in B taken for pathway analysis in g:Profiler. Bubbles are coloured based on each clusters and size is proportional to number of genes in specific GO term. D. Venn diagram showing integrative analysis of significant GOs in most variable methylated (Figure 2D) and concordant (Figure 2E) genes (left panel). Bubbles plot showing common pathways between genes upregulated/hypomethylated and in clusters 3 and 4 (middle panel) or genes downregulated/hypermethylated and in clusters 1 and 2 (left panel). Bubbles are coloured based on FDR and size is proportional to number of genes of specific GO term.

**Supplementary Figure 3.**
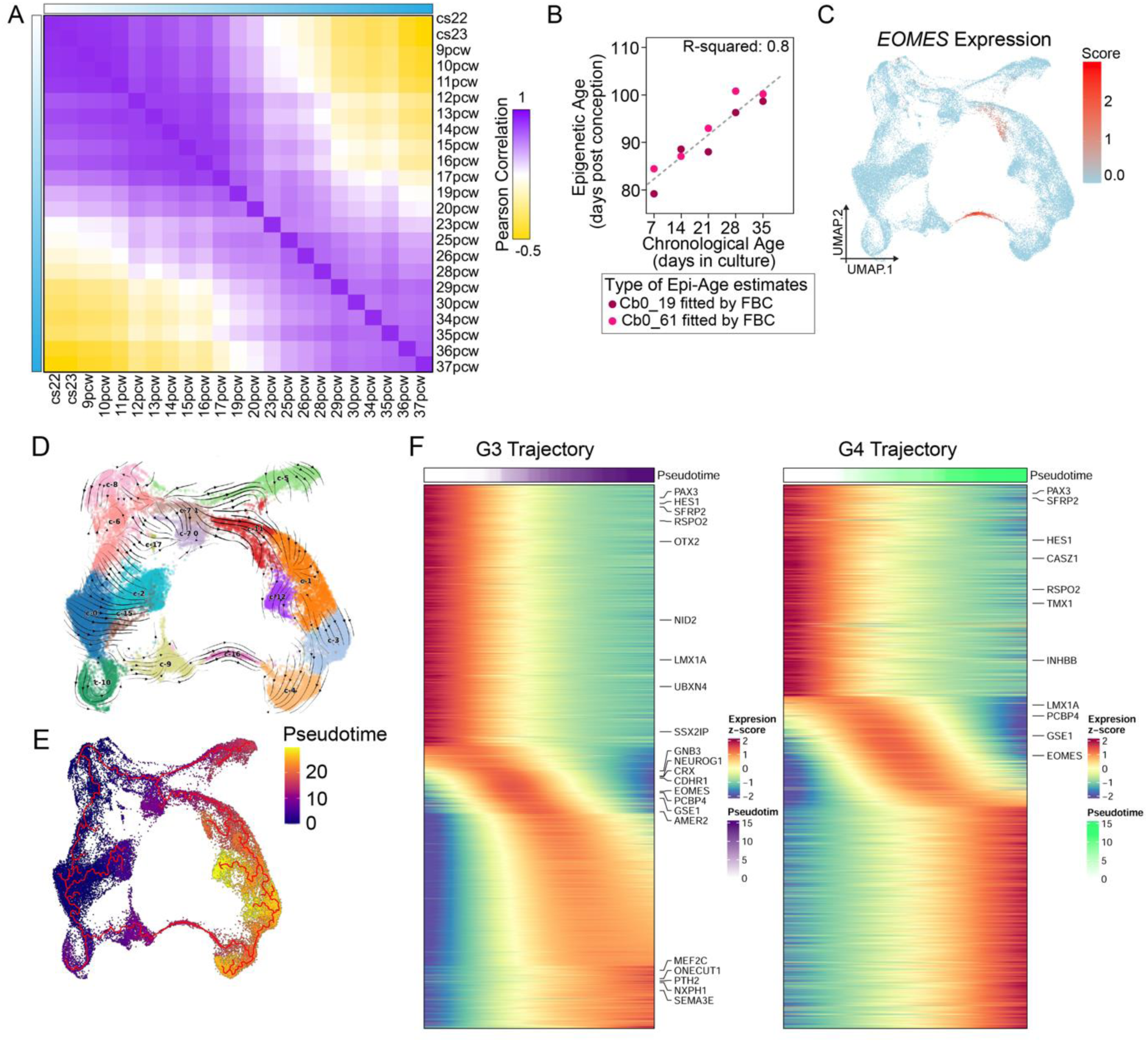
CbO clusters harbouring MB cell-of-origin are enriched for distinct G3 and G4 MB associated gene signatures. A. Heatmap showing correlation of methylation status of top 1% age-dependent probes between human developing cerebellum samples, demonstrating a linear correlation through development. B. Correlation plot between days of maturation of CbO_19 and CbO_61 (chronological age) and the modelled epigenetic ages from the foetal brain clock^5^. C. Uniform Manifold Approximation and projection (UMAP) of Day 35/49 CbO showing alra imputed expression score for *EOMES*. D. Day 35/49 CbO UMAP with RNA velocity streams shown, calculated in velocyto and scVelo. E. Pseudotime analysis of Day 35/49 CbO UMAP calculated in Monocle3, with cycling progenitors cluster (see Figure 1G) selected as the root of pseudotime. F. Plots of z-scored gene expression through pseudotime within G3 and G4 trajectories defined in Figure 3K.

**Supplementary Figure 4.**
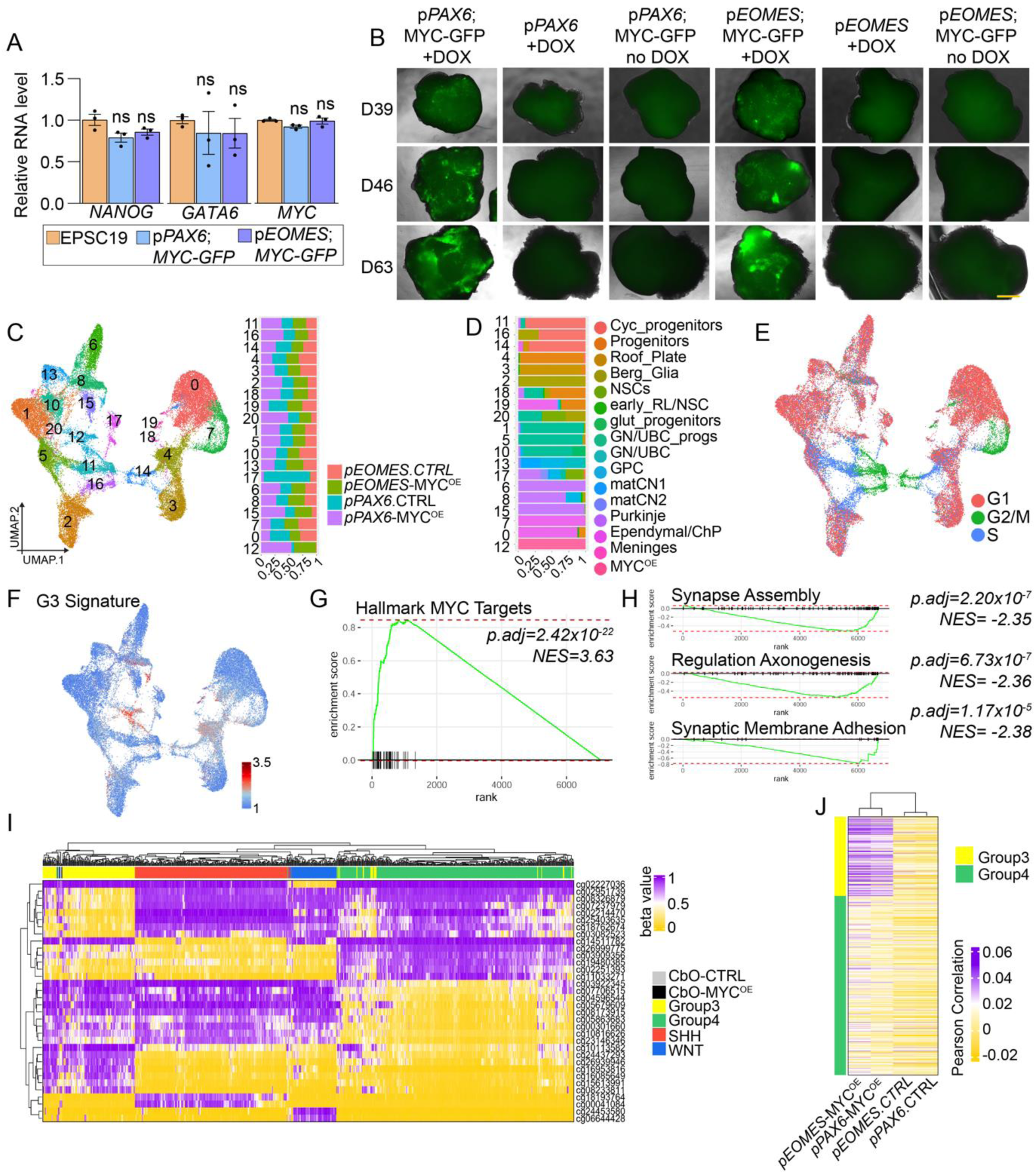
Cerebellar organoid editing strategy displays specificity to medulloblastoma lineages-of-origin and similar methylation profiles. A. qPCR data of naïve expanded potential stem cells (EPSC19) and EPSC19 infected with *pPAX6;MYC-GFP* or *pEOMES;MYC-GFP* editing constructs outlined in Figure 4A for pluripotency (*NANOG)* and differentiation (*GATA6)* markers as well as c-*MYC.* Relative RNA level signifies dCT values expressed relative to naïve EPSC19 (2^-ddCT). ns = not significant, one-way ANOVA. n = 3 technical replicates. B. Representative images of CbO19 at Day 36, 46 and 63 of culture infected with different combinations of *pPAX6, pEOMES* or *MYC-GFP* editing constructs (refer to Figure 4A), with or without the addition of doxycycline (DOX) to organoid media from Day 35 of culture. C. UMAP plot of scRNA-Seq data from D63 *pEOMES;MYC-GFP, pPAX6;MYC-GFP* (*pEOMES*-MYC^OE^, *pPAX6*-MYC^OE^) and control (CTRL) CbOs coloured by Louvain clusters (left). Barplot showing the relative proportion of cells clusters in each sample (right). D. Barplot showing the relative proportion of cell types in each cluster in **C** (labels transferred from D35/49 control CbOs (Figure 1G). E. UMAP plot shown in C, coloured by cell cycle phase. F. UMAP plot shown in C, showing Group 3 MB signature score^6^. G. Gene set enrichment plot of the HALLMARKs MYC TARGETS V1 gene set in the MYC^OE^ cluster markers (Figure 4C), NES=3.63, padj=2.24E-22. H. Gene set enrichment plot showing depletion of gene sets associated with neural differentiation in MYC^OE^ cells relative to D35/49 G3/4 lineage-of-origin cells. I. Clustered heatmap of medulloblastoma (MB) samples taken from Cavalli et al.^7^ and edited CbO samples based on their beta values of probes shown to delineate MB subgroups^8^. J. Clustered heatmap of edited CbO sample correlation to group 3 and group 4 MBs from the Cavalli et al. dataset^7^ on the beta values of differentially methylated probes between *MYC* overexpressing (*pEOMES*-MYC^OE^, *pPAX6*-MYC^OE^) and control medulloblastoma lineage-of-origin samples (*pEOMES.*ctrl, *pPAX6.*ctrl).

**Supplementary Figure 5.**
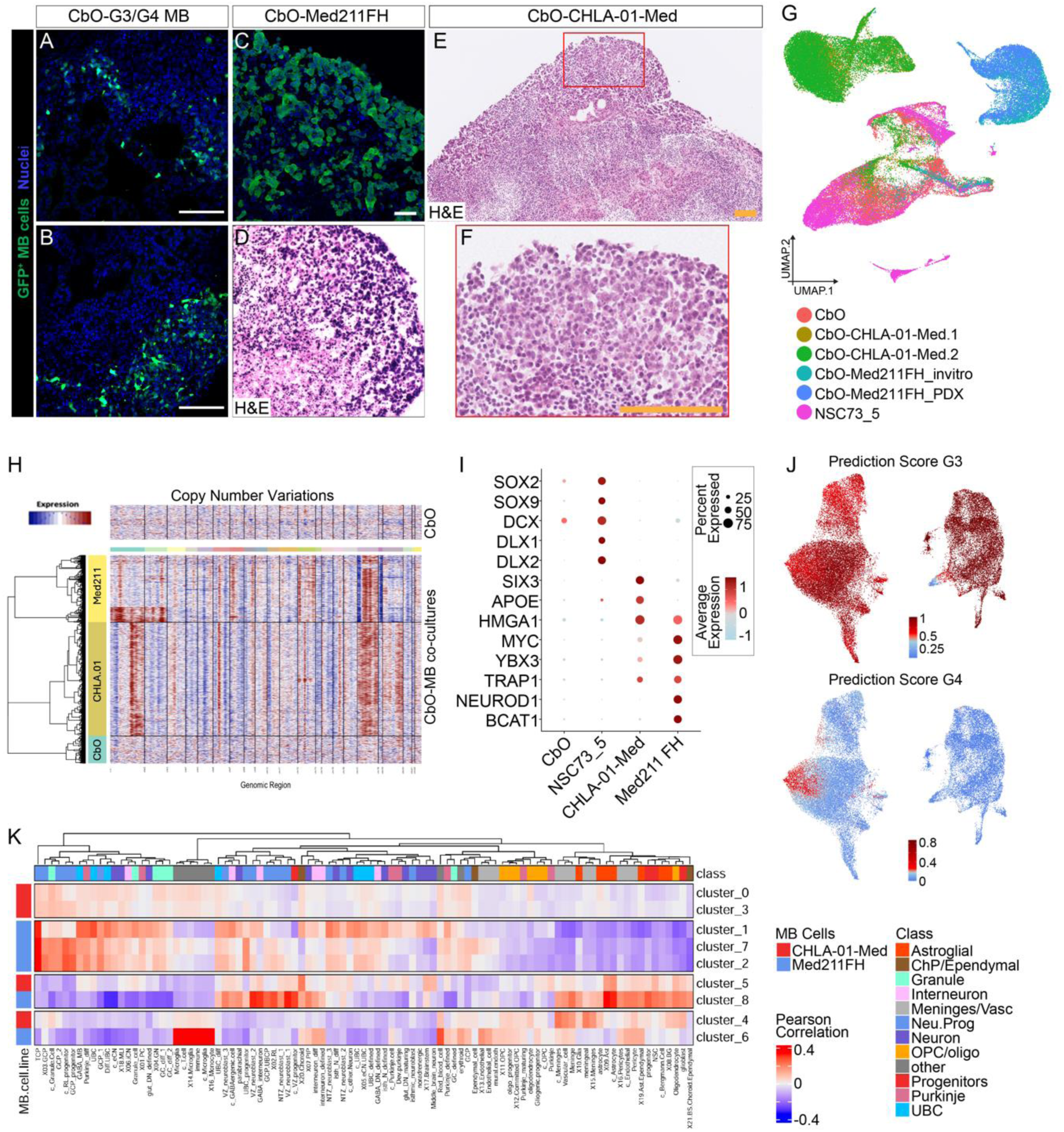
Medulloblastoma (MB) cells display the morphology of human MB upon co-culture with cerebellar organoids (CbO) and retain their genetic features. A. Representative immunofluorescence staining of GFP^+^ ICb1299 G3/4 MB cells upon fourteen days of co-culture with CbO_61, scale bar = 100 µm. B. Representative immunofluorescence staining of GFP^+^ CHLA-01R-Med recurrent G3/4 MB cells upon fourteen days of co-culture with CbO_61, scale bar = 100 µm. C. Representative immunofluorescence staining of GFP^+^ Med211FH cells upon fourteen days of co-culture with CbO_61, scale bar = 50 µm. D. H&E staining of CbO-Med211FH co-cultures upon 14 days of co-culture with MB cells, scale bars = 50 µm. E. H&E staining of CbO-CHLA-01-Med co-cultures upon 14 days of co-culture with MB cells, scale bars = 100 µm. F. H&E staining of CbO-Med211FH co-cultures upon 14 days of co-culture with MB cells, scale bars = 100 µm. G. UMAP plot of scRNA-Seq data from D49 control CbO_61, CbO-Med211-FH, CbO-CHLA-01-Med and control CbO-NSC73.5 co-cultures, coloured by sample. H. Hierarchical clustering of inferred copy number profiles separates co-cultured CHLA-01-Med (suede) and CbO-Med211-FH (yellow) from CbO (cyan) cells. I. Dot plot showing key marker genes that distinguish co-culture NSC73_5, CHLA-01-Med and CbO-Med211-FH cell populations from CbO. J. UMAP plot of scRNA-Seq data from MB cells only from D49 CbO-Med211-FH and CbO-CHLA-01-Med CbO-MB co-cultures, coloured by label transfer scores for Group3 (top), Group 4 (bottom). K. Heatmap showing correlation of co-cultured Med211-FH and CHLA-01-Med scRNA-Seq clusters to reference cell type from four independent scRNA-seq datasets of the human developing foetal cerebellum^1–4^.

**Supplementary Figure 6.**
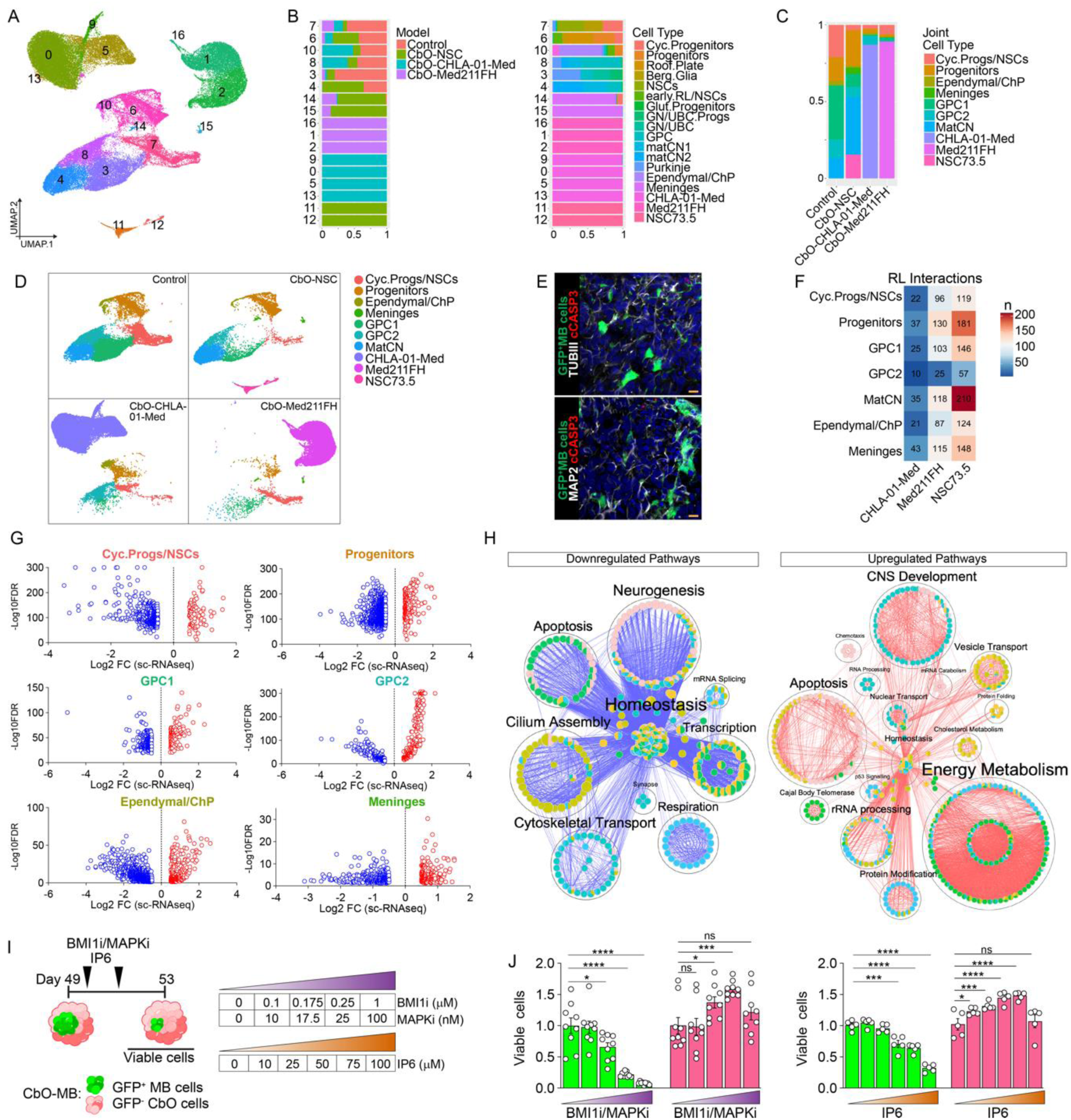
Co-culture with medulloblastoma cells modulates transcriptional profiles of specific cerebellar organoid lineages. A. UMAP plot of scRNA-Seq data from D49 control CbO_61, CbO-Med211-FH, CbO-CHLA-01-Med and control CbO-NSC73_5 co-cultures, coloured by Louvain clusters. B. Barplot showing proportion of each cluster made up of cells from CbO_61 control, CbO-Med211-FH, CbO-CHLA-01-Med or CbO-NSC73_5 co-culture samples (left). Barplot showing proportion of each cluster made up of different cell types from CbO_61 control, CbO-Med211-FH, CbO-CHLA-01-Med or CbO-NSC73_5 co-culture samples (right). For CbO clusters cell labels were computationally transferred from D35/49 controls (Fig 1G). C. Barplot showing proportion of samples in A, B made up by the joint cell type labels. D. Uniform manifold approximation and projections (UMAPs) plot of scRNA-Seq data from D49 control CbO, CbO-Med211-FH, CbO-CHLA-01-Med and control CbO-NSC73_5 co-cultures. Cells are coloured by their assigned cell type and UMAPs are split according to experimental model. E. Immunohistochemistry images of cerebellar organoids co-cultured with NSC73.5 cells for 14 days. Upper panel: GFP (green), cCASP3 (red) and TubIII (white); Lower panel: GFP (green), cCASP3 (red) and MAP2 (white). Scale bars = 10 µm. F. Heatmap showing the total number of predicted significant receptor-ligand interactions (regardless of directionality) between co-cultured CbO-Med211-FH, CbO-CHLA-01-Med and control CbO-NSC73_5 cells and CbO_61 cell populations. G. Volcano plots showing significant differentially expressed genes (DEGs) upon MB cells co-culture in indicated CbO_61 cell populations. Red and blue dots represent significantly upregulated and downregulated genes respectively. H. Bubble plot showing top 100 significant (adjusted p-value < 0.05) GO Biological Processes and Molecular Functions of up- (red) and down- (blue) regulated genes from 6 volcano plots shown in G taken for pathway analysis in g:Profiler. Bubbles are coloured based on each cell population and size is proportional to number of genes in specific GO term. I. Schematic of CbO-CHLA-01-Med treatment with tables reporting the concentrations of PTC209 (BMI1i), PD329501 (MAPKi) and IP6 used. J. Viability assays of GFP^+^ MB cells (green) and GFP^−^ CbO cells (pink) in CbO-MB upon 4 days of treatment with increasing concentrations of BMI1 and MAPK inhibitors (left) or IP6 (right) as shown in Figure 5H and 5I respectively. All graphs report mean ± SEM, number of independent CbO analyzed is reported in each graph, one-way ANOVA ^✱^p<0.05, ^✱✱✱^p<0.001, ^✱✱✱✱^p<0.0001, ns = not significant.

